# The Interferon Resistance of Transmitted HIV-1 is a Consequence of Enhanced Replicative Fitness

**DOI:** 10.1101/2021.12.18.473292

**Authors:** Elena Sugrue, Arthur Wickenhagen, Nardus Mollentze, Muhamad Afiq Aziz, Vattipally B Sreenu, Sven Truxa, Lily Tong, Ana da Silva Filipe, David L Robertson, Joseph Hughes, Suzannah J Rihn, Sam J Wilson

**Author notes:** These authors contributed equally to this work (Elena Sugrue, Arthur Wickenhagen).

## Abstract

HIV-1 transmission via sexual exposure is a relatively inefficient process. When successful transmission does occur, newly infected individuals are colonized by either a single or a very small number of establishing virion(s). These transmitted founder (TF) viruses are more interferon (IFN) resistant than chronic control (CC) viruses present 6 months after transmission. To identify the specific molecular defences that make CC viruses more susceptible to the IFN-induced ‘antiviral state’ than TF viruses, we established a pair of fluorescent GFP-IRES-Nef TF and CC viruses and used arrayed interferon-stimulated gene (ISG) expression screening. The relatively uniform ISG resistance of transmitted HIV-1 directed us to investigate the underlying mechanism. Our subsequent *in silico* simulations, modelling, and *in vitro* characterisation of a model TF/CC pair (closely matched in replicative fitness), revealed that small differences in replicative growth rates can explain the broad IFN resistance displayed by transmitted HIV-1. We propose that the apparent IFN resistance of transmitted HIV-1 is a consequence of enhanced replicative fitness, as opposed to specific resistance to individual IFN-induced defences.

## INTRODUCTION

The type I interferon (IFN) response is one of the earliest immune defences deployed against invading pathogens, including HIV-1 [1]. IFN signalling results in the expression of hundreds of IFN-stimulated genes (ISGs), many of which restrict virus replication, thereby creating an antiviral state [2, 3].

HIV-1 sexual transmission is a surprisingly inefficient process [4], with >98% of sexual exposure events not resulting in transmission [5]. Therefore, the virions in fluids from infected individuals are usually unable to establish a productive infection in a new host. In the unusual event of successful transmission, there is typically a severe genetic bottleneck, such that infection is typically established by just a single genetic variant (or a very small number of variants), described as the transmitted founder (TF) virus(es) [6-10]. Such limited transmission arises from both physical and immunological barriers that restrict viruses from the typically large and diverse population present in the donor from productively infecting target cells in a new host [11-13]. Thus restrictive transmission can be described as a stochastic process, where small fitness advantages can lead to successful infection [14].

Different phenotypic TF virus properties have been proposed to contribute to successful infection, and their relative importance in the context of a stochastic process considered [14, 15]. Subtle differences in envelope glycoprotein structure between TF viruses compared to chronic control (CC) viruses present six months after infection have been observed [16], and hypothesized to potentially be favoured at transmission [17, 18]. The differential use of CCR5 receptors between TF and CC viruses has also been noted [19]. TF viruses have been proposed to often be more IFN resistant than viruses isolated during chronic infection [20-22]. Given their often proposed reduced sensitivity to IFN, TF viruses have fully functional accessory genes (*vpu, vif* and *nef*) that can counteract ISGs [23-25], and these genes are maintained over the course of infection [26]. However, the relative importance of these phenotypic properties for successful infection has also been debated, with some describing the process as substantially stochastic [27, 28].

Two theories predominate when considering the transmission bottleneck and the phenotypic properties of TF viruses. The first is that TF viruses are phenotypically unique and more resistant to the IFN response [20, 22, 29], the second is that TF viruses are simply more fit in terms of replicative success [14, 30]. TF viruses have previously been observed to be apparently relatively IFN resistant and uniquely resistant to specific ISGs, such as the IFITMs, with this resistance reported to decrease during chronic infection and prolonged exposure to the host immune response [31]. Recent work characterising 500 clonally derived HIV-1 isolates confirmed the dynamic nature of IFN resistance during chronic infection, and showed how the relative contribution of IFN to HIV-1 control varies at different stages of infection, or after interruption of antiviral therapy [21]. Other studies into the importance of viral replicative capacity on HIV-1 immunopathogenesis have also revealed its importance in the TF virus, independent of host protective genes and viral load [30, 32].

While there is growing evidence describing the dynamic nature of the IFN response and HIV-1 control, and how TF viruses are more resistant to the resultant antiviral state [21], mechanistic understanding about the specific molecular defences that make CC viruses more susceptible to the antiviral state is currently incomplete. Here, following our initial endeavour to identify the specific antiviral defenses resisted by TF viruses, we reveal the relatively uniform ISG resistance profile of a representative transmitted HIV-1. Our subsequent *in silico* simulations, modelling, and *in vitro* characterisation of a model TF/CC pair (closely matched in replicative fitness), demonstrates that small differences in replicative growth rates can explain the broad IFN resistance displayed by transmitted HIV-1. These unanticipated observations suggest that small fitness advantages can underlie apparent IFN resistance. Importantly, these data suggest that the ‘replicative success’ vs ‘IFN resistance’ theories of successful HIV-1 transmission are not in opposition, but are instead inherently linked.

## RESULTS

### Transmitted HIV-1 is more resistant to IFN

Identifying specific molecular defences that explain the relative resistance of HIV-1 transmitted founder (TF) viruses to IFN, when compared to matched chronic control (CC) viruses present 6 months after transmission, first required selection of an appropriate matched TF/CC pair for screening experiments. To select a pair, we examined the replication of four previously described infectious molecular clone (IMC) pairs [20] in immortalized human T cells over several days. To start the initial infections, which all used equivalent 0.01 multiplicities of infection (MOIs), we used virions pseudotyped with the vesicular stomatitis virus glycoprotein (VSV-G) in order to circumvent the low levels of infection typically observed with some IMCs. To visualise the spread of the unmodified viruses in subsequent rounds of infection, we used an LTR-GFP reporter cell line, MT4 TMZR5 cells [33], which fluoresce green when infected and could be monitored daily via flow cytometry. Notably, three of the pairs tested (CH040, CH236 and CH850) exhibited large differences in replicative fitness in the absence of IFN, which would make examining the relative resistance of these pairs to IFN challenging (Fig 1A-B). In contrast, the CH058 TF/CC pair exhibited similar replicative kinetics in the absence of IFN, as well as high levels of overall infection (Fig 1A-B). Therefore, the CH058 pair was selected for use as a model pair for our subsequent screening experiments. Additionally, to remove any confounding issues from pseudotyping with VSV-G, the CH058 IMC pair was additionally propagated in TMZR5 cells (using stocks produced without pseudotyping) to generate sequence-verified working stocks for subsequent experiments.

**Figure 1.**
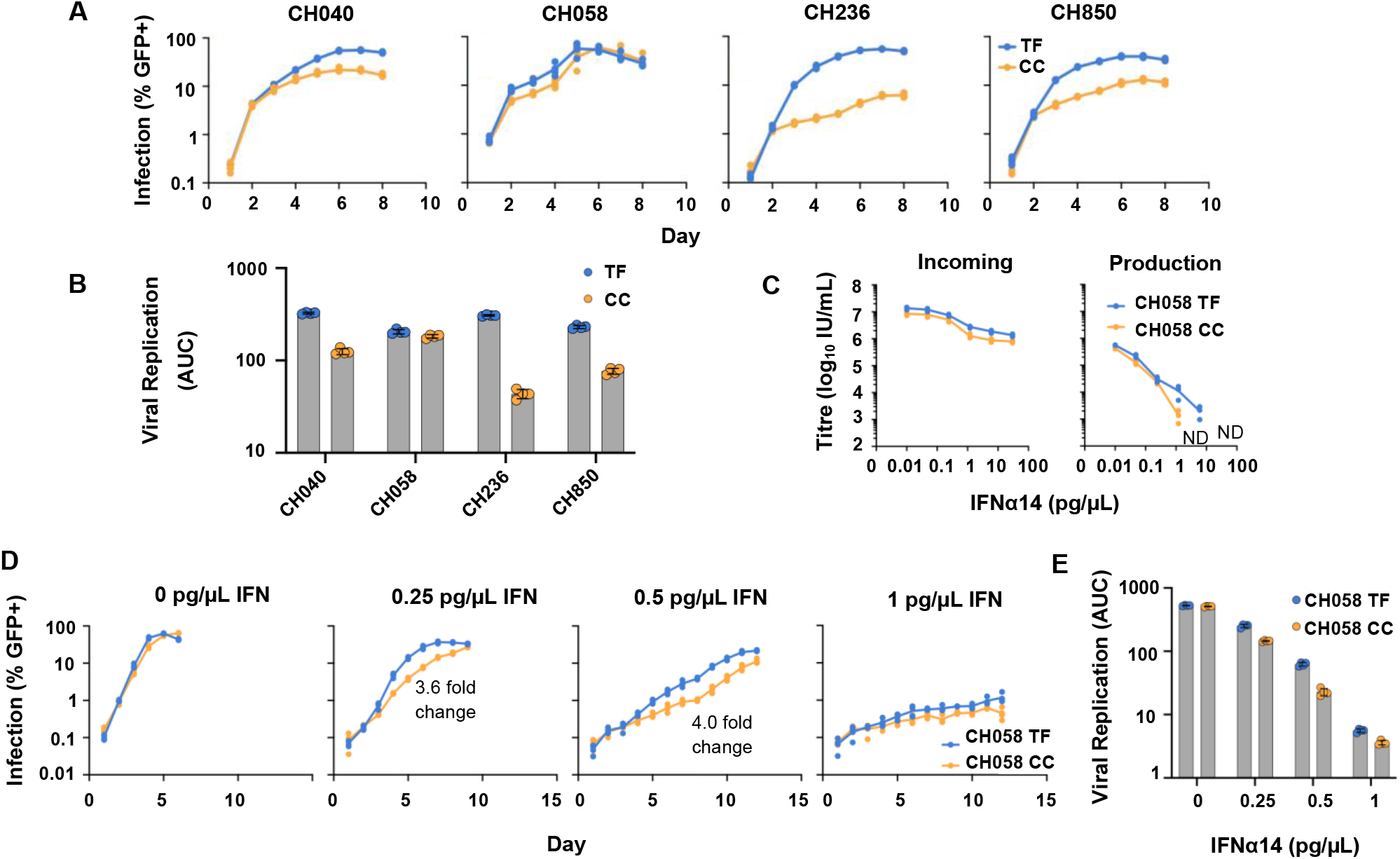
The CH058 TF virus is more resistant to IFN than its matched CC virus. (A-B) TMZR5 GFP-reporter cells were challenged with matched TF and CC VSV-G pseudotyped IMC pairs and sampled daily to monitor virus spread. GFP-positive cells were enumerated via flow cytometry. (C) To investigate the effect of IFN early in the viral life cycle (incoming infection), TMZR5 cells were pre-stimulated with different doses of IFNα14 for 24 hours prior to infection with serially diluted CH058 TF and CC viruses. To limit replication to a single cycle, infected cells were treated with dextran sulphate 17-18 hours post infection. The percentage of GFP positive cells was determined at 48 hours post infection using flow cytometry. To investigate the effect of IFN late in the viral life cycle (production effect), TMZR5 cells were pre-treated with IFNα14, and after 24 hours cells were challenged with CH058 TF and CC viruses at an MOI of 0.5 for 6 hours, before the inoculum was removed and washed with phosphate buffered saline (PBS). At 46-48 hours post infection, cell-free, filtered virus containing supernatants were titrated on TMZR5 cells. ND indicates not detected. (D-E) TMZR5s were treated with the indicated dose of IFNα14 for 24 hours before being challenged with the CH058 TF and CC virus pair. Cells were sampled daily to monitor virus spread and GFP-positive cells were enumerated via flow cytometry. Annotated fold change values refer to the maximum difference in (%) infection out of the timepoints tested. Viral spreading replication experiments took place on two occasions and a typical result is Shown.

ISGs induced by type I IFNs can confer protection against HIV-1 during early (incoming) infection [34-38] and during late (production) of infectious progeny [24, 39]. Pre-treatment of TMZR5 cells with varying concentrations of IFN*α*14 stimulated modest ∼10-fold protection against incoming infection from the CH058 TF and CC viruses tested (Fig 1C). At doses lower than 0.24 pg/μl, the incoming titres of both TF and CC viruses were unaffected. As the concentration of IFN*α*14 increased above 0.24 pg/μl, the infectivity of both TF and CC viruses was moderately suppressed. We subsequently determined the infectious yields of CH058 TF and CC viruses using TMZR5 cells stimulated with varying concentrations of IFN*α*14 (Fig 1C). Without IFN stimulation, both viruses displayed a similar level of infectious progeny virions. Elevating the dose of IFN*α*14 caused substantial reduction in the infectious production of both CH058 viruses. Strikingly, at 6.0 pg/μl and higher, the infectious yield of the CH058 CC virus was reduced to below the level of detection, whereas infectious CH058 TF was readily detectable (Fig 1C). This indicates that IFN*α*14 caused a stronger reduction in the infectious yield of the CC virus than of the TF virus (using the CH058 pair), and also suggests that IFN*α*14 confers a relatively weak early block (∼10-fold) and potent late block (>200-fold) to HIV-1 CH058 in TMZR5 cells.

To further investigate the impact of IFN*α*14 on CH058 replication, we examined the ongoing replication (over a longer timescale) of the CH058 pair in cells pre-treated with a range of IFN*α*14 doses. Notably, the TF virus again outperformed the CC virus across all the IFN doses tested, despite comparable replication kinetics in the absence of IFN (Fig 1D-E). Because of the proapoptotic effect of IFNs, the viability of IFN-treated cells was also assessed in parallel cultures. The majority of IFN doses tested exhibited a live population of 80-90%, with the highest dose tested (1 pg/ µL) displaying a ∼60% live population (S1).

### ISG expression screening reveals multiple ISGs that inhibit the CC virus more potently than transmitted HIV-1

We have previously used arrayed ISG expression screening to identify antiviral factors targeting a range of viruses [2, 40, 41]. Although HIV-1 has previously undergone large-scale ISG and CRISPR screening [2, 3, 42] a matched TF/CC pair has not yet been investigated in this way, and could reveal specific molecular defences resisted by transmitted HIV-1. We therefore conducted ISG screening using our human ISG library, which includes >500 unique ISGs encoded in SCRPSY lentiviral vectors (Fig 2A), in conjunction with a GFP-encoding TF/CC pair (CH058) we developed in order to enable easy quantification of virus infection using flow cytometry. To construct this GFP TF/CC pair, we inserted an IRES-GFP cassette between *env* and *nef*, to create the viruses we will refer to as the CH058 GIN (GFP-IRES-*nef*) viruses (Fig 2A).

**Figure 2.**
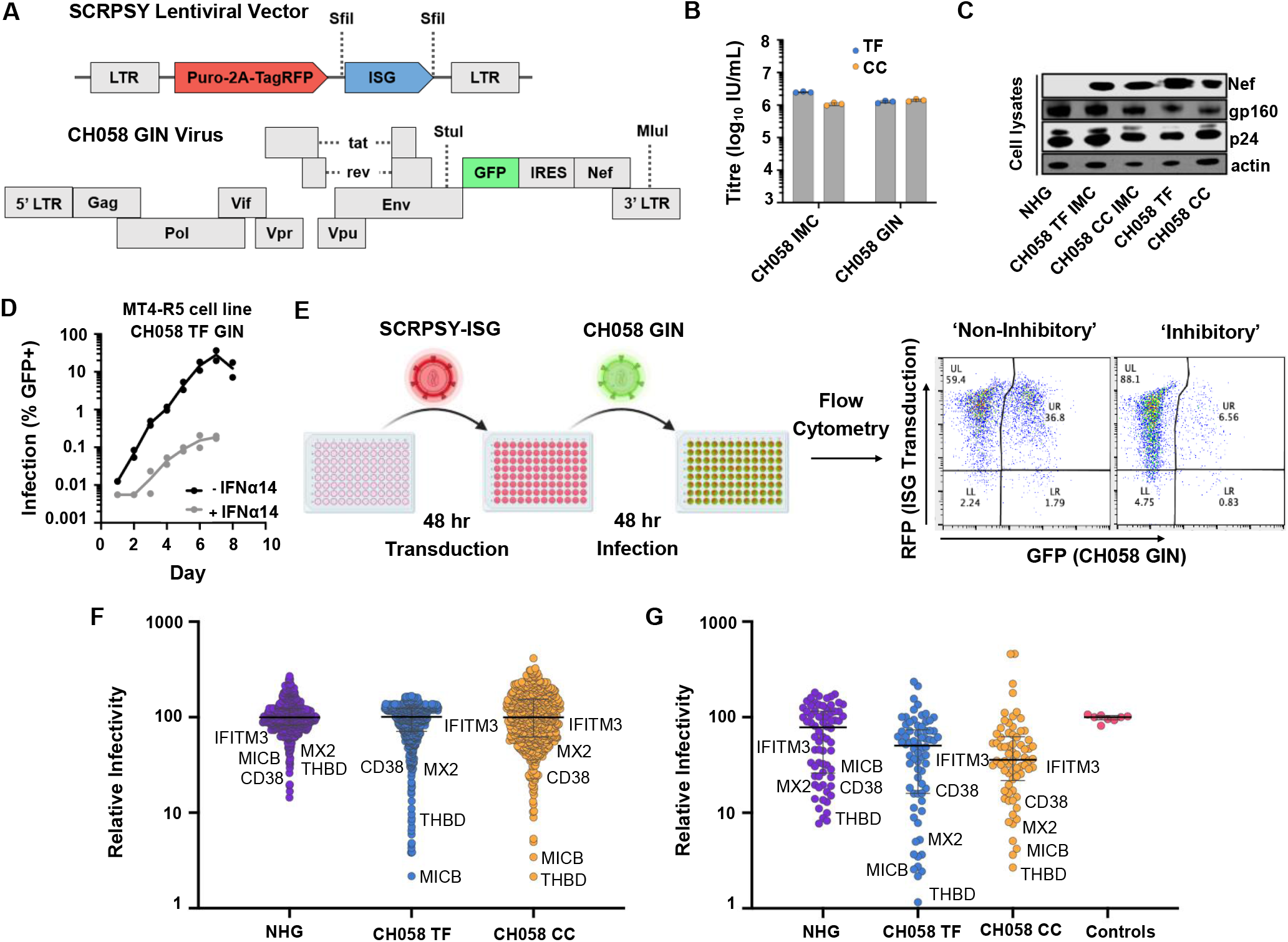
Arrayed ISG expression screening identifies multiple ISGs that inhibit transmitted HIV-1 and its matched CC virus. (A) A schematic of the SCRPSY lentiviral vector (GenBank accession KT368137.1) used to deliver ISGs, and of the fluorescent GFP-IRES-Nef (GIN) TF and CC viruses used. (B) The infectious titres of CH058 IMC and CH058 GIN viruses in a single cycle of infection in TMZR5 cells. Cells were treated with dextran sulphate 17-18 hours post infection, the percentage of GFP positive cells was determined at 48 hours post infection using flow cytometry. (C) Western blot analysis of viral antigens presents in TMZR5 cell lysates infected with indicated virus for 48 hours. (D) Growth kinetics of CH058 TF GIN in clonal MT4 CCR5-R126N in the absence and presence of 0.5 pg/μl IFNα14. Cells were treated with IFNα14 24h prior to infection. (E) A schematic of the ISG screening pipeline used in panel F. (F) Normalized infection (median centred) of cells expressing different ISGs (each dot represents the observed infection in the presence of a single ISG). (G) Validation screen, conducted as in E and F, of ISGs ‘hits’ selected that were more inhibitory than human Mx2 in panel F. Empty SCRPSY transduced MT4-R5 cells were used as controls.

As described above, the CH058 pair was chosen as an ideal pair for screening because they exhibit the most similar replication kinetics (Fig 1A-B). Stocks of the GIN viruses were prepared (Fig 2B) and viral protein expression (Fig 2C) was assessed and was found to be comparable to the unmodified CH058 IMCs. We then elected to conduct the ISG screens in MT4 cells modified to express a signalling-defective variant of CCR5 [43]. Importantly, these cells, referred to as MT4-R5 cells, are both readily transduced by our ISG library, and also support efficient HIV-1 replication that is potently inhibited by type I IFN treatment (Fig 2D). We transduced the MT4-R5 cells with the ISG-encoding lentiviral library and, 48 h later, infected these cells with CH058 TF GIN and CH058 CC GIN viruses (Fig 2E). At 96 h post-infection, the level of CH058 GIN infection in the presence of each individual ISG was quantified using flow cytometry (Fig 2E-F).

Due to the low levels of infection that would occur in a single replication cycle from the GIN variants of CH058, we assessed multi-cycle infection in the ISG screens for these viruses. However, as these multi-cycle infections could mask potential anti-HIV-1 genes acting early in the life cycle, we also conducted a single-cycle ISG screen using lab-adapted HIV-1 NHG (Fig 2F), which is an NL4.3-derived virus, that contains portions of HxB2 envelope, and that encodes GFP in place of *nef* [44]. Following completion of these screens, and in order to pinpoint specific ISGs that inhibit HIV-1, we identified all genes that showed equivalent or stronger inhibition than the known anti-HIV-1 ISG Mx2 in any individual screen [45]. We then subtracted known IFNβ/ISRE-inducing ISGs [2] from this list and re-examined the ability of independent lentiviral vector preparations encoding each of these potentially antiretroviral ISGs to inhibit HIV-1 (Fig 2G). Following this ‘miniscreen’, we selected all the ISGs that exhibited inhibition equivalent or stronger than displayed by IFITM3, an ISG resisted by transmitted HIV-1 [31], (25 candidate genes) for subsequent analysis.

We next examined the ability of these 25 candidate anti-HIV-1 effectors and a vector control to inhibit CH058 TF GIN and CH058 CC GIN HIV-1 in a multi-cycle infection on MT4-R5 cells transduced with a new batch of lentiviral vectors expressing the candidate effectors (Fig 3A). To potentially exclude genes from our final selection that are either ISRE- or cell death-inducing, we conducted four subtractive screens on the gene list from Fig 2G including our 25 candidate effector genes. We tested the ability of all genes to induce cell death in MT4 or TMZR5 cells (Fig 3B), tested the cell viability of MT4 cells transduced with these genes (Fig 3B) and assessed ISRE stimulation in a MT4-ISRE-GFP cell line transduced with these genes (Fig 3C). Genes showing more than 2.1-fold increase in any of these screens were excluded from further analysis. Additionally, we used published studies from the interferome v2.0 database [46] to investigate the ‘ISG-ness’, or degree to which a gene is stimulated by interferon (Fig 3D). This led us to exclude AKT3, FAM134B and THBD, as their type I IFN stimulation profile showed downregulation in more than half the published datasets where differential expression was observed (Fig 3D). Based on their strong anti-HIV-1 activity (Fig 3A), no considerable induction of cell death or ISRE stimulation (Fig 3B-C), and strong IFN-stimulation (Fig 3D), we selected CD38, CD80, FNDC3B, IFITM3, MICB, Mx2, SCARB2 and TMEM140 as the final 8 genes which exhibited strong anti-HIV-1 activity in our screens. Mx2 and IFITM3 are ISGs known to target HIV-1 [31, 35, 36, 45, 47, 48] whereas the other genes have not yet been intensively investigated with regards to anti-HIV-1 activity.

**Figure 3.**
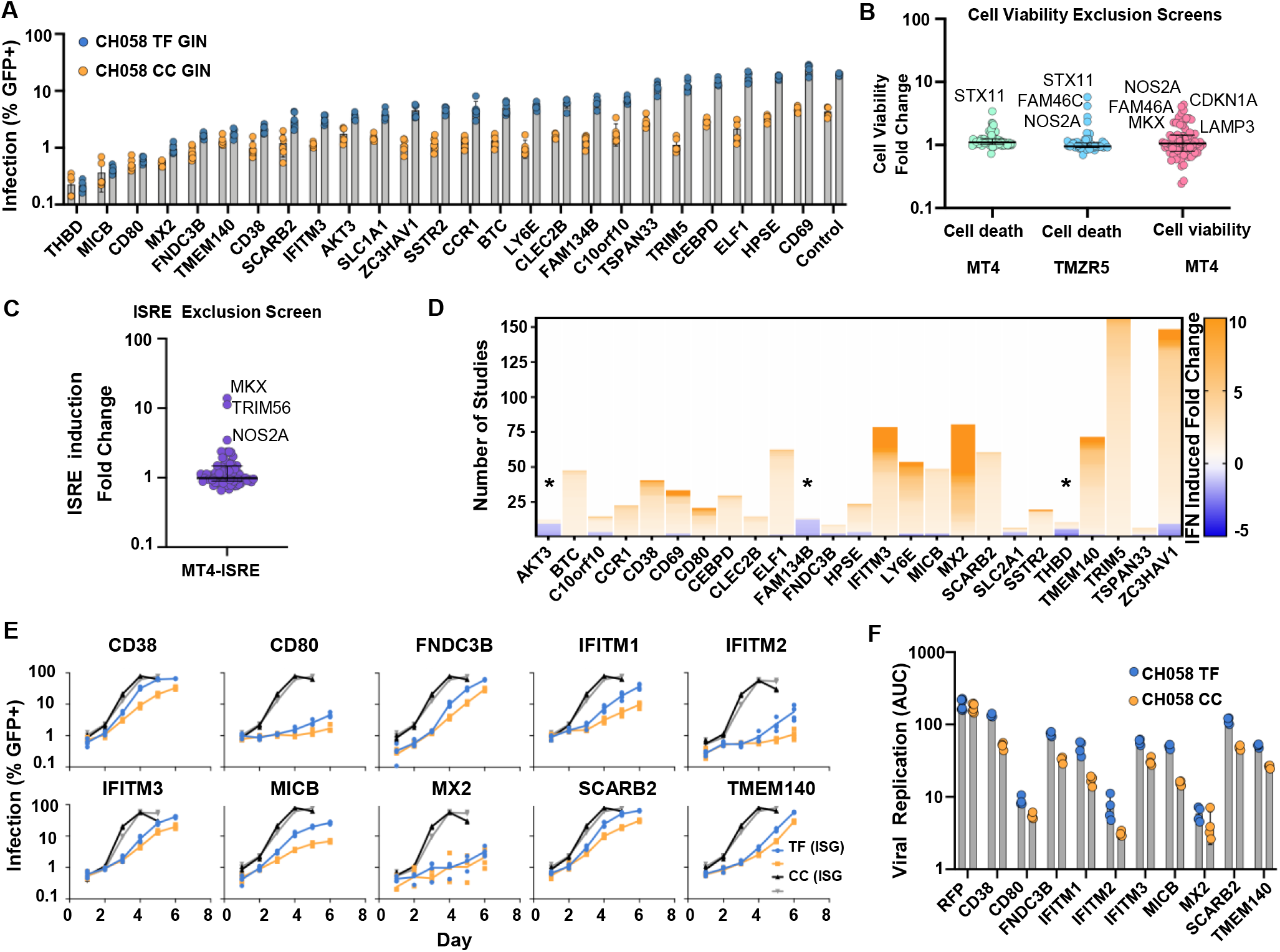
Anti-HIV-1 ISGs inhibit the CC virus more potently than transmitted HIV-1. (A) Candidate anti-HIV-1 effectors that were more inhibitory than the known anti-HIV-1 effector IFITM3 were tested against CH058 GIN TF and CC viruses on MT4-R5 cells, as in 2F-G. (B-C) ISGs in Fig 3A were tested for ability to induce cell death or ISRE. In B, MT4 and TMZR5 cells were tested for cell death when expressing ISGs by flow cytometry using the LIVE/DEAD fixable dead cell stain kit (Invitrogen) and MT4 cell viability was additionally tested using the luminescence based CytoTox-Glo Cytotoxicity assay (Promega). In C, IFN induction by candidate ISGs was measured by flow cytometry using MT4 cells expressing an ISRE-GFP construct. (D) Candidate ISGs from Fig 3A were checked for their changes in expression upon type I IFN stimulation using the interferome database to determine their ‘ISG-ness’. (E-F) Validation of the 8 most potent anti-HIV-1 effector ISGs, along with IFITM1 and IFITM2 controls, against CH058 TF and CC viruses, In E, TMZR5 cells transduced with pLV constructs containing the indicated ISGs or RFP as a control, were challenged with CH058 TF or CC and sampled daily to monitor virus spread. GFP-positive cells were enumerated using flow cytometry. Viral spreading replication experiments took place on two occasions, a typical result with contemporaneous controls is shown. In F, data from panel E represented as area under the curve (AUC).

The final candidate anti-HIV-1 effectors, alongside IFITM1 and IFITM2 controls, were then subcloned into a pLV lentiviral expression vector, which subsequently allowed stably modified GFP-reporter TMZR5 cells [33] expressing each ISG to be established. These cells were infected with a low MOI (0.01) using non-modified CH058 TF and CH058 CC virus stocks and sampled daily to monitor virus spread (Fig 3E-F). All 10 exogenously expressed genes robustly inhibited HIV-1 replication when compared to an RFP control. Yet strikingly, comparisons of the CC and TF CH058 virus results revealed that the transmitted variant of CH058 was relatively resistant to all the ISGs tested except Mx2.

Given that six of the genes identified using our pipeline (CD38, CD80, FNDC3B, MICB, SCARB2 and TMEM140) have not been characterised as encoding anti-HIV-1 effectors, we wanted to further investigate the role endogenous expression of these ISGs could play in the anti-HIV-1 effects of IFN. We thus used western blots to screen the endogenous expression levels of all six ISGs in a variety of cell lines and primary cells, in the presence and absence of IFN, in order to detect IFN-induced expression, and to also identify the best targets for CRISPR/Cas9 manipulations (S2). Analysis of these western blots identified both CD38 and SCARB2 as potential endogenous effectors, as both exhibited readily detectable endogenous expression, with observable increases in the presence of IFN. In contrast, the endogenous expression of CD80, FNDC3B and MICB was only weakly IFN inducible, and levels were considerably lower than the exogenous levels that inhibited HIV-1 in Fig 3E (S2). In addition, we were unable to convincingly detect TMEM140 expression.

To investigate whether endogenous SCARB2 and CD38 might inhibit HIV-1, we disrupted these loci using CRISPR/Cas9. We examined the protein expression of each target using transduced ‘bulk’ populations and identified guides that reduced CD38 expression in PM1 cells, as well as guides that attenuated SCARB2 expression in TMZR5 cells (S2). We followed HIV-1 replication in these two cell lines with the greatest reduction in endogenous expression (of CD38 or SCARB2), and observed no notable changes in HIV-1 replication compared to the non-targeting control cell lines (in the presence and absence of IFN). This evidence suggests CD38 and SCARB2 are unlikely to play a major role in the inhibition of HIV-1 by type I IFNs *in vivo* (S2).

### Small differences in either growth rate between a virus pair, or in resistance to inhibition, are amplified by logistic growth

The observation that transmitted HIV-1 was relatively resistant to multiple ISGs, including ISGs whose endogenous expression we did not find to be inhibitory, led us to hypothesize that the apparent difference in IFN sensitivity between TF and CC viruses was not driven by resistance to specific antiviral defences, but was instead a consequence of different virus growth rates of the TF and CC viruses (i.e., differences in replicative fitness as opposed to genetic resistance to specific effectors with anti-HIV-1 activity). We therefore simulated viral growth curves under these competing hypotheses, assuming logistic growth with a starting population of 100 infected cells and a carrying capacity of 10000 cells (broadly matching conditions in our experiments). In one set of simulations, viruses differed in growth rate only, with the second virus having a lower growth rate than the first. While the inhibition scaling factor was fixed at 1, giving both viruses equal sensitivity to the growth rate inhibitor (which could be IFN). In the second set of simulations, the underlying growth rates of both viruses were equal, but the second virus was more sensitive to the growth rate inhibitor (mimicking scenarios where one virus is more sensitive to specific antiviral effectors). Interestingly, these simulations revealed that a small difference in growth rate was sufficient to recapitulate the apparent IFN resistance observed in our experimental data (Fig 4A; c.f. Fig 1A, D & Fig 3E).

**Figure 4.**
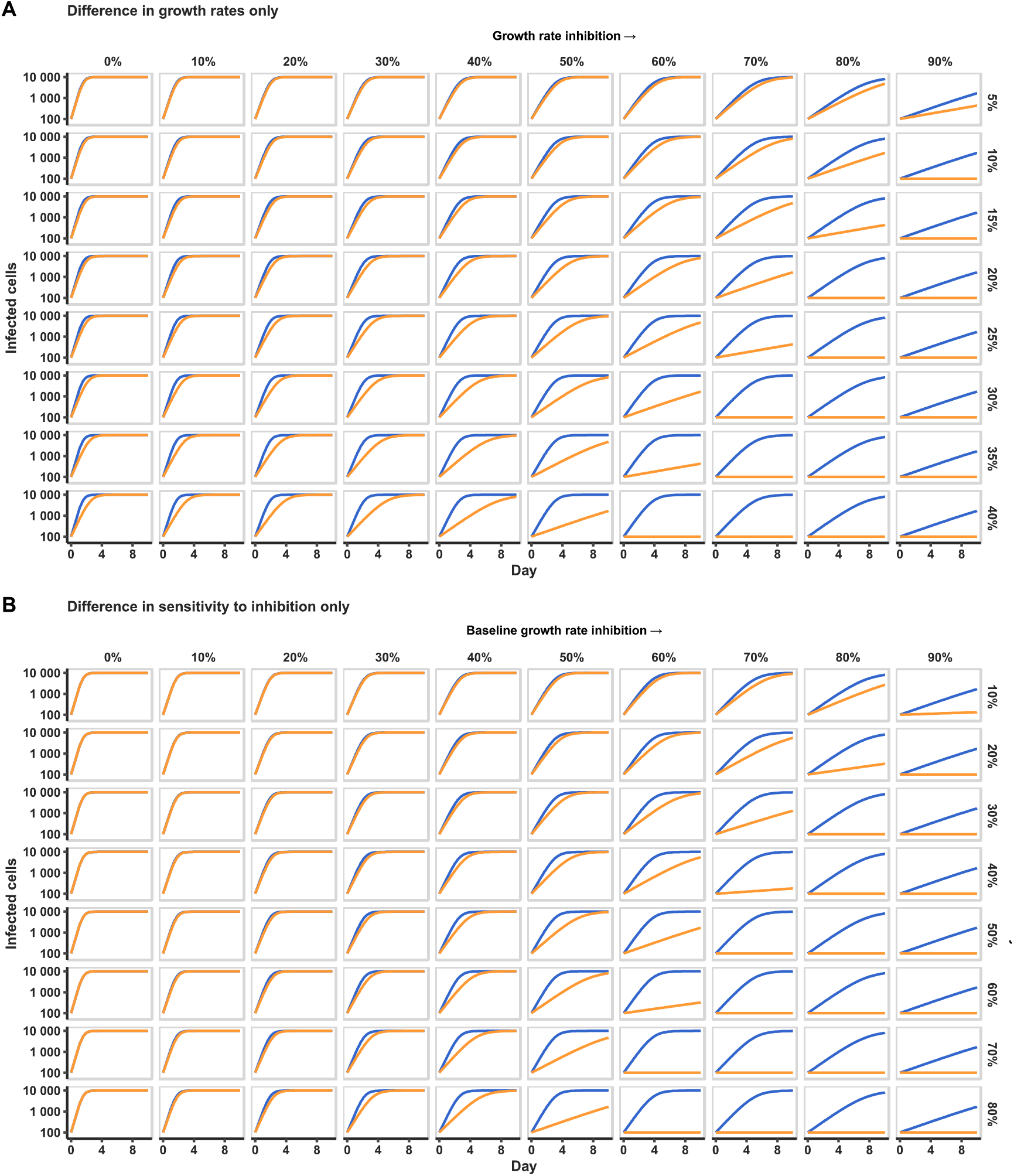
Both small differences in growth rate between a virus pair, and differences in sensitivity to growth inhibition, can explain relative interferon resistance of transmitted HIV-1. (A) Logistic growth simulation of two viruses, where the growth rate of virus two (orange) is scaled relative to that of virus one (blue). Rows represent an increasing difference in growth rates between viruses (i.e., an increasing difference in replicative fitness). Both viruses experience the same relative growth rate inhibition (columns) mimicking increasing IFN stimulation. (B) Simulations in which growth rates are identical, but virus two (orange) is more sensitive to the growth rate-inhibiting factor (i.e., one virus is more sensitive to antiviral effectors). Columns represent an increasing growth rate inhibition (mimicking increasing IFN stimulation), while rows represent an increasing difference in the sensitivity of the two viruses to that inhibition.

When a sufficiently large growth rate impediment (mimicking the antiviral state induced by IFN (c.f. Fig 1A)) was applied to the simulated growth curves, the slower-growing virus was undetectable, whereas the faster growing virus grew exponentially and overwhelmed the culture (Fig 4A-B). Under normal culture conditions in the absence of IFN (using MOIs typical of these experiments), the lag phase is largely by-passed due to the rapid growth rate of HIV-1. Inhibiting the growth rate (simulating IFN stimulation), slowed growth to the point where the lag phase became observable (Fig 4A, S3) Notably, even a 10% difference in growth rates led to the fitter virus infecting ∼10-fold more cells by day 4 in the presence of high concentrations of IFN (Fig 4A; c.f Fig 1A). Alternatively, similar dynamics could be produced by assuming identical growth rates but a difference in inhibitor sensitivity (mimicking IFN sensitivity, specifically the CC virus being more sensitive to specific ISGs), as has been hypothesised before [29] (Fig 4B). Similar results were obtained when simulating exponential growth (unpublished observations). However, the logistic growth assumed for this work more closely matches the replication of HIV-1 observed than exponential growth, and has previously been used to explain the dynamics of HIV-1 replication [49, 50].

### Transmitted HIV-1 has a higher growth rate, independent of interaction with IFN

We next wanted to use these simulations as a basis to further determine whether the observed TF/CC growth kinetics reflect differences in TF/CC viral growth rates or differences in their sensitivity to inhibition. To do this, we implemented a viral spreading assay using the CH058 TF/CC pair over a more expansive range of IFN*α*14 doses, with a focus on increments between 0 and 0.5 pg/µL, as this is where the largest difference in replication was observed (Fig 1D). TMZR5 cells were pre-treated with IFN for 24 hours prior to virus inoculation, and the infection levels were monitored daily via flow cytometry (Fig 5A). We next fitted two alternative logistic growth models to the observed number of infected (GFP+) cells (Fig 5B-C). These models incorporated regressions on the growth rate and carrying capacity parameters (with the latter used to account for IFN toxicity, which had the effect of reducing the number of cells available to infection at higher doses of IFN). In the differential sensitivity model (Fig 5B), differences in the IFN-sensitivity of the CC virus were allowed, whereas, in the constant sensitivity model (Fig 5C), additional sensitivity of the CC virus was not considered (i.e. fixed at zero). Importantly, both models maintained the patterns seen in our experimental observations, and a model assuming no difference in the effect of IFN on growth rates between viruses (‘constant sensitivity’, Fig 5C) fitted the experimental data just as well as one allowing for ‘differential sensitivity’ (ΔAIC = 1.81; Fig 5B). Indeed, both models utilised reduced baseline growth rate of the CC virus to achieve optimal fitting (Fig 5D) and increased IFN-sensitivity of the CC virus was not required for optimal fitting (Fig 5D-F). Importantly, the difference between the fitted growth rates of TF and CC viruses did not increase over a range of IFN doses, indicating that neither virus was substantially more or less sensitive to IFN (Fig 5E). Instead, we found the constant ∼17% lower growth rate of the CC virus (95% confidence interval: 12.1 – 18.8% lower) (Fig 5D-E), was sufficient to recapitulate the apparent IFN resistance of transmitted HIV-1. Thus, modest increases in the replicative fitness of TF viruses underlie the interferon resistant phenotype, and could be crucial for breaking through the bottleneck of HIV-1 transmission.

**Figure 5.**
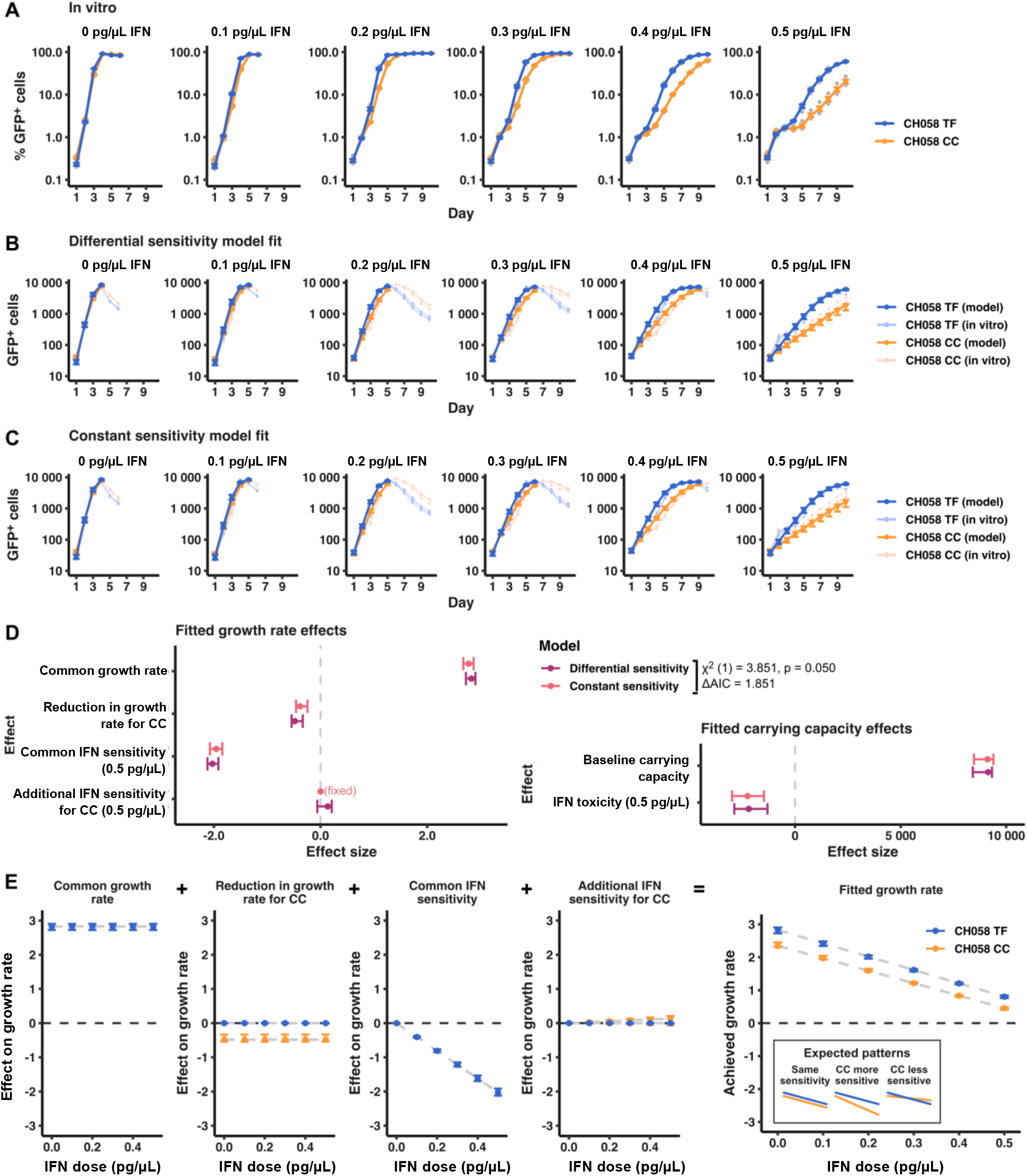
Modelling of virus growth curves indicates that CH058 CC has a lower growth rate than its TF progenitor but is not otherwise more sensitive to IFN. (A) TMZR5 cells were pre-stimulated with different doses of IFNα14 for 24 hours prior to infection with CH058 TF or CC virus, and sampled daily to monitor virus spread. GFP-positive cells were enumerated via flow cytometry. Blue and orange points show means (+/-standard error) across 4 experimental replicates, while translucent points show individual observations. Viral spreading replication experiments took place on two occasions, a typical result is shown. (B) Model fit for the differential sensitivity model, in which viruses are allowed to vary in both baseline growth rate (i.e., growth rate in the absence of interferon) and in their sensitivity to interferon. (C) Model fit for the constant sensitivity model, in which viruses differ only in their baseline growth rate. In both B and C, opaque points show predictions from the fitted model, along with 95% confidence intervals. Translucent points and lines underneath show the observed (experimental) replicate growth curves also present in A. For each treatment, only the initial timepoints until at least one replicate curve decreased by >30% relative to its preceding observation were used in model fitting (i.e. points after HIV-1 had overwhelmed the culture were not considered). (D) Effect sizes for both differential and constant sensitivity models (maximum likelihood estimates and 95% confidence intervals). Model fit was compared by likelihood ratio test and AIC, with results shown in the legend. (E) Illustration of effects making up the fitted growth rates in the differential sensitivity model. Effects were modelled as additive, allowing us to separate discrete contributions to the growth rates needed to recapitulate in vitro data as illustrated in B. Points show maximum likelihood estimates, while error bars show 95% confidence intervals. An inset in the final panel (right) illustrates expected patterns under different hypothesized scenarios; in particular, if the CC virus was more sensitive to IFN, its growth rate would have diverged from that of the TF virus at increasing IFN doses.

### Transmitted HIV-1 is more resistant to both IFN and antiretroviral compounds

If the apparent increased IFN resistance of transmitted HIV-1 is simply a by-product of enhanced replicative fitness, we hypothesized that transmitted HIV-1 would also be more resistant to other inhibitory agents, including those not normally encountered during sexual transmission. We therefore investigated whether a transmitted HIV-1 was more resistant to antiretroviral compounds. Crucially, we selected two antiretroviral compounds that would target viral proteins that were identical in the model TF and CC virus pair. There are only 8 amino acid differences between the TF and CC CH058 IMCs (*gag*: G251E, *tat*: K29R, *env*: T232A, N338D, R579S, A830T, *rev*: R54Q, *nef*: G113E, [20]) and the viruses encode identical protease and reverse transcriptase (RT) enzymes. Therefore, we considered the ability of the RT inhibitor azidothymidine (AZT), and the protease inhibitor nelfinavir (NFV), to inhibit TF and CC CH058. Strikingly, the TF was relatively resistant to both AZT and NFV (Fig 6 A-D), reminiscent of the resistance to IFN exhibited in Figure 1 and Figure 5. Given the absence of sequence diversity in the protease and RT of the TF and CC viruses, this experiment strongly suggests that enhanced replicative fitness underlies the apparent resistance of this transmitted HIV-1 to antiretroviral compounds.

**Figure 6.**
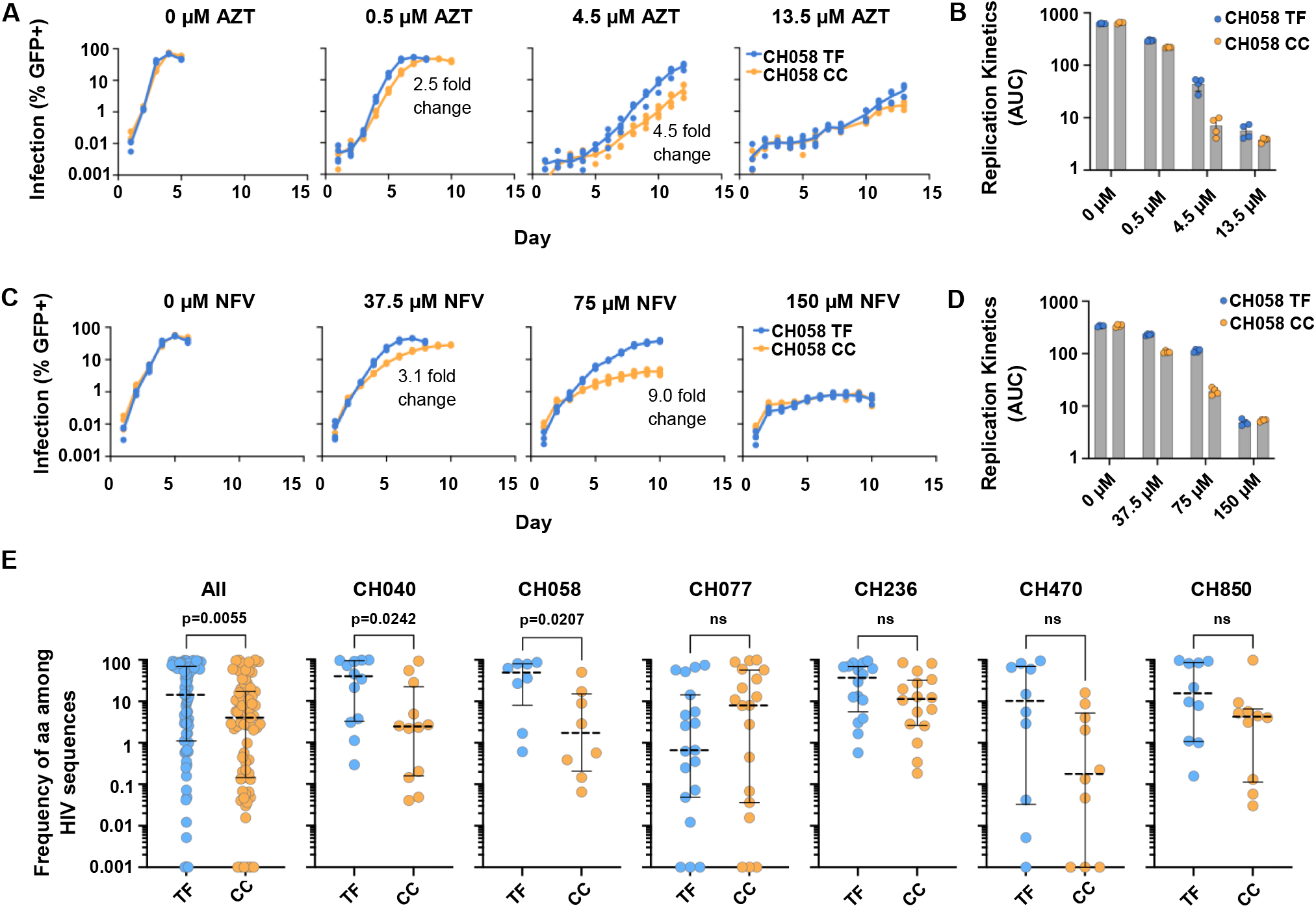
Transmitted HIV-1 is more resistant to antiretroviral drugs and tends to access more prevalent sequence space than matched chronic viruses. (A-B) TMZR5 cells were pre-treated with a range of azidothymidine (AZT) doses for 2 hours prior to infection with the CH058 TF or CC virus, cells were sampled daily to monitor virus spread. GFP positive cells were quantified via flow cytometry. (C-D) TMZR5 cells were pre-treated with a range of nelfinavir (NFV) doses and infected, sampled and quantified as in A. Viral spreading replication experiments took place on two occasions, a typical result is shown. (E) Amino acid substitution (or variation) frequencies for sites that exhibit amino acid differences between the matched TF/CC pairs are shown using 4568 sequences for tat to 19237 for nef in subtype B and from 1548 sequence in rev to 4345 in env for subtype C from the Los Alamos sequence database. Each point represents one of the amino acid sites that differs between the TF and CC in that virus pair. The amino acid frequencies of the equivalent sites in HxB2 are also shown as a comparator. Significance was determined using a Mann-Whitney test.

### TF viruses overwhelmingly exhibit prevalent residues at polymorphic sites

A selection bias favouring transmission associated founder effects of viruses encoding amino acids associated with increased replicative fitness has previously been identified [14], and additional HIV-1 studies have also shown that amino acid prevalence and fitness can be closely linked [51-53]. To carefully characterise any sequence changes between the TF and CC viruses investigated in this work (CH040, CH058, CH236 and CH850), we sequenced these pairs via Illumina MiSeq. The sequenced IMCs had 100% coverage with a minimum mean depth of 5229 (Fig S4). Two additional pairs investigated in a foundational paper describing the resistance of TF viruses to IFN (CH077 and CH470) were included in our analysis for completeness [20].

This sequencing allowed us to identify the amino acid sites that exhibit changes between the matched TF/CC pairs (‘divergent TF/CC sites’) and allowed us to compare amino acid frequencies at these divergent sites to a reference HIV-1 sequence, HxB2 (S5). In order to consider the frequencies of amino acids at divergent TF/CC sites in the context of a more global representation of HIV-1 sequences and of genome evolution, we next obtained all available sequences of HIV-1 subtype B and subtype C from the Los Alamos HIV sequence database (www.hiv.lanl.gov/). Strikingly, when we evaluated amino acid usage of HIV-1 at the divergent TF/CC sites for all Los Alamos sequences (of the relevant subtype), we found that the TF viruses tended to utilise residues that were more frequently accessed by HIV-1, while the CC viruses tended to use residues that were used less frequently (p=0.0055) (Fig 6E, S5). This trend of TF viruses accessing more frequently utilised sequence space at the divergent TF/CC positions seems likely to be a consequence of these residues conferring increased replicative fitness. We speculate that this trend could be due to constraints that are absent in a new host (such as acquired immune attack or antiretroviral therapy), selecting for transmitted variants that access optimal sequence space for replication in a naïve host. We used CH058 to investigate how many sites of change were associated with immune escape using the HIV mutation browser (https://hivmut.org/) [54]. We found that seven out of eight sites had publications associated with drug or immune escape (*gag* 248 [55, 56], *env* 232 [57], 339 [58, 59], 588 [60], 747 [61], *rev* 54 [62], and *nef* 108 [63, 64]). Interestingly, amongst the virus pairs tested, CH077 contrasts from this observed trend, as the distribution of conserved frequencies appears lower for the TF than the CC CH077 virus. CH077 is also observed to be an outlier in a recent work investigating the fitness of TF/CC pairs [32]. In that study, no significant fitness difference was detected from a single passage competitive fitness assay between the TF and the CC virus, and a difference in fitness could only be determined by passaging the mixture of cell-free viruses three times [32].

## DISCUSSION

The propensity of TF viruses to be IFN-resistant has previously been identified, and is often described as an important determinant of successful HIV-1 transmission [22, 29, 65]. Over the course of chronic HIV-1 infection, and on-going accumulation of diversity, variants with altered properties that can evade multiple host defences (such as neutralizing antibodies) arise [66-69]. However, despite this differential susceptibility to host defences, paired TF/CC viruses have not previously been subjected to arrayed ISG expression screening. Our aim was to identify specific molecular defences that make CC viruses more susceptible to the IFN-induced antiviral state than TF viruses. Through our screening we identified multiple ISGs that could inhibit both our reporter TF and CC viruses. Remarkably, for the majority of ISGs tested, the CC virus was more sensitive to ISG-mediated inhibition than the TF virus. Thus, the consistent ISG resistance exhibited by transmitted HIV-1 hinted at a single, common, underlying mechanism.

Both relative resistance to specific antiviral defences and improved replicative fitness have been described in other work as characteristics of TF viruses when compared to CC viruses [21, 22, 30, 31]. We compared both characteristics and demonstrated that small differences in modelled growth rate between a virus pair, or differences in inhibition strength (mimicking IFN treatment), were amplified by logistic growth, and these closely matched the patterns seen in our experimental data. Our subsequent statistical modelling clearly indicated that a difference in IFN sensitivity was not the cause of the relative IFN resistance of transmitted HIV-1. Instead, a minor difference in replicative fitness explained the observed IFN resistance of transmitted HIV-1. Indeed, the idea that reduced replicative fitness can mechanistically underly increased IFN sensitivity has previously been proposed as a general process that could tip the balance in favour of the host and influence virus pathogenesis and host range [70-72].

The relatively small (17%) difference in growth rate between TF and CC CH058 we calculated from the modelling was intriguing. Previous work investigating the genetic fragility of HIV-1 capsid revealed that the majority of amino acid substitutions in capsid caused a greater fitness reduction (70% of random single amino acid changes in capsid caused at least a 50% reduction in fitness [73]). Additionally, research into the fitness landscape of HIV-1 Gag revealed that making multiple sequence changes that were predicted to impact fitness (based on sequence prevalence) to Gag protein also resulted in replicative capacity differences much greater than our observed difference [52]. However, single unfavourable amino acid changes or multiple changes that were predicted to not greatly impact fitness had replicative differences in a similar range to ours. Additional work investigating HIV-1 escape mutations showed that substitutions in genes demonstrating slow evolution resulted in dramatic losses in replicative fitness, that are again much greater than our observed changes, while substitutions in HIV-1 genes with rapid evolution did not have a negative impact on pathogen fitness [74]. These data indicate that while relatively small differences in growth rate, similar to the CH058 pair, may appear very small, they can still be phenotypically significant.

To further clarify the role of replicative fitness compared to IFN/inhibitor sensitivity in TF vs CC viruses, we designed an experiment centred around sensitivity to inhibitors that target enzymes (protease and reverse transcriptase) that have identical sequences in the matched CH058 TF/CC pair. The striking observation that the CH058 TF was more resistant to AZT and NFV than a matched CC virus (Fig 6), despite having identical RT and protease sequences, clearly implicated the underlying role of enhanced replicative fitness amongst transmitted variants. It correspondingly seems likely that other TF/CC phenotypes may also be explained by replicative fitness [75]. In particular, Hertoghs *et al*. showed that when other matched HIV-1 TF/CC pairs were studied in Langerhans cells (LCs), a mucosal macrophage subset that has been shown to have a protective role in HIV-1 transmission, most of the TF viruses tested were able to infect LCs, whereas matched CC viruses had lost this ability [75]. Additionally, research into hepatitis C virus (HCV) has also shown that high replicative fitness is linked to increased resistance to antiviral agents [76, 77] and resistance to lethal mutagenesis [78]. Importantly, as we chose to focus our experimental work on the TF/CC pair whose fitness was most closely matched (Fig 1A), it is likely that our results underemphasise the role that enhanced replicative fitness of the TF virus may have in other contexts. This is particularly true given that in most pairs, the fitness difference could be easily detected without any additional inhibitory agent, indicating that the difference would likely be greatly exaggerated by IFN treatment.

While we examined a single model TF/CC pair using reporter cell lines, recent work testing ten matched TF/CC pairs in competitive fitness assays in primary cells supports our conclusions [32]. In the work of Wang *et al*., ten CC viruses were less fit than their matched TF viruses, suggesting our conclusions could be broadly applicable [32]. While TF viruses may not always be the most fit or apparently most IFN-resistant compared to non-transmitted viruses from a transmitting partner, our study reinforces the notion that TF viruses have higher replicative fitness than comparable chronic viruses [21], and we propose that replicative fitness underlies the apparent IFN resistance of transmitted HIV-1. To this end, we also found that at polymorphic sites, transmitted HIV-1 tended to utilise more frequently accessed sequence space (that is therefore likely to have higher replicative fitness) than CC viruses. Additionally, that many divergent sites between a representative pair are associated with drug/immune escape. Thus, our work is consistent with the idea that acquired immune responses increasingly drive chronic HIV-1 into a constrained sequence space that is resistant to immune attack but less replicatively fit (in the absence of immune attack). During transmission to a naïve host, the now less fit in this new context (but immune resistant) variants are outcompeted by their fitter, immune sensitive, counterparts (that perhaps originate from a reservoir established early after infection). Thus, the observable relative IFN resistance of transmitted HIV-1 can be achieved through enhanced replicative fitness, as opposed to resistance to specific antiviral effectors. Notably, a nonspecific mechanism does not downplay the importance of IFN resistance as a key phenotypic property of transmitted HIV-1. Moreover, nonspecific IFN resistance in no way depreciates the pivotal role that IFN responses likely play as a barrier to HIV-1 transmission [79].

## MATERIALS AND METHODS

### Cells

Adherent HEK 293T cells were propagated from lab stocks maintained in Dulbecco’s modified Eagle’s medium (DMEM) supplemented with 10% fetal calf serum (FCS) and 10 μg/ml gentamicin. Suspension MT4 cells were expanded from lab stocks and maintained in RPMI medium supplemented with 10% FCS and 10 μg/ml gentamicin. MT4-LTR-GFP indicator cells (TMZR5 cells) have been modified to express the CCR5 receptor and contain a cassette in which hrGFP expression is driven by the HIV-1 LTR and have been described previously [2, 33]. The MT4 CCR5-R126N cells, referred to as MT4-R5 cells in this work, were produced through the PCR of genomic DNA extracted from TMZR5 cells to generate the R126N CCR5 product, and to also introduce SfiI restriction sites at the 5’ and 3’ ends of CCR5 gene, enabling cloning into an MLV-based vector (primer pair AA-099-LPCX CCR5-F 5’-CTCTCTGGCCGAGAGGGCCATGGATTATCAAG TGTCAAGTCCAATC-3’ and AA-100-LPCX CCR5-RC 5’-TCTCTCGGCCAGAGAGGCCTCACAAGCCCACA GATATTTCCTGC-3’). Following transduction of MT4 cells, a limited dilution strategy was implemented to select a cell line that fostered replication and maintained IFN sensitivity. All lentivirus transduced cells were selected and cultured in medium additionally supplemented with 2 μg/ml puromycin (Melford Laboratories), 5 μg/ml blasticidin (Melford Laboratories) or 1 mg/ml G418 (Invitrogen) as appropriate.

### Retroviral vectors and plasmids

SCRPSY (KT368137.1) lentiviral vector has been previously described [2], pLV-EF1a-IRES-Neo (Addgene plasmid #85139) was modified to include SfiI sites flanking the transgene ORF by inserting the TagRFP (or gene of interest) ORF with flanking SfiI sites between the unique BamHI and EcoRI restriction sites using PCR (primer pair AW177-BamHI-SfiI-RFP-F’ 5’-CTCTCGGATCCGGCCGAGAGGGCCATGAGCGAGCTGATTAAG-3’ and AW178-EcoRI-SfiI-RFP-R’ 5’-CTCTCGAATTCGGCCAGAGAGGCCTCACTTGTGCC CCAG-3’). Gene editing was achieved using the lentiCRISPRv2-Blast system [80].

### Replication competent viruses

Lentivirus stocks (Table 1) were generated through transient transfection of HEK 293T cells in the presence/absence of pCMV-VSV-G using polyethylenimine (PEI). The following clones were used: replication-competent GFP-encoding pNHG (JQ585717) [44, 81]. A panel of full-length transmitted/founder (TF) and matched chronic control (CC) HIV-1 infectious molecular clones were obtained as generous gifts from Beatrice Hahn and Stuart Neil. In all cases, supernatants were harvested at ∼48 h post transfection and clarified using a 0.45-μm-pore-size filter and stored at −80°C. CH058 working stocks were additionally propagated for 10 days in TMZR5 cells after transfection.

**TABLE 1:** Viruses used in this study

| Virus (subtype) and designation | Description | Accession number | Co-receptor usage | Reference(s) |
| --- | --- | --- | --- | --- |
| HIV-1 Group M (B) |  |  |  |  |
| NHG | GFP in place of *nef* | JQ585717 | X4 | [44] |
| CH058-GIN | TF CC GFP and IRES inserted between *env* and *nef* genes | MW535546<br>MW535544 | R5<br>R5 | This Study |
| CH040 | TF CC | MW535542<br>MW535541 | R5<br>R5 | [82] |
| CH058 | TF CC | MW535545<br>MW535543 | R5<br>R5 | [82] |
| HIV-1 Group M (C) |  |  |  |  |
| CH236 | TF CC | MW535549<br>MW535548 | R5<br>R5 | [20] |
| CH850 | TF CC | MW535552<br>MW535551 | R5<br>R5 | [20] |

### IFNα14 production and quantification

Stat1 deficient U3A fibroblasts, a generous gift from Stephen Goodbourn, were utilised to minimise the presence of secreted ISGs in IFN preparations. These U3A cells, which lack STAT1, were modified to produce IFN under a doxycycline-inducible system. In order to efficiently generate high quantities of IFNα14, engineered U3A cells expressing IFNα14 were seeded into 10-cm dishes at a ratio of 1:3 to achieve maximum confluency prior to stimulation with 125 ng/ml of doxycycline (DOX). The DOX treated cells were incubated for 24 hours to allow sufficient expression of IFNα14 before the supernatants were harvested and purified using a 0.45-μm filter. The biological units of recombinant human IFNα14 produced in this study were determined using ISRE-GFP expressing HEK293T cells. Cell-free supernatants containing IFNα14 were 1.5-fold serially diluted and titrated onto 2.0 × 10^5^ cells/ml of ISRE-GFP cells in a 96-well plate. Titration of IFNα14 was carried out in parallel with commercial IFN, where commercial IFN stocks were used to generate a standard curve for a dose determination. Based on the calculation, the estimated concentration of IFNα14 was 1153.2 pg/μl. To assess the toxicity of IFNα14 treatment the LIVE/DEAD fixable green dead cell stain kit (Invitrogen) was used.

### Arrayed ISG expression screening

The ISG overexpression screening was completed similarly to previously described protocols [2, 40]. In short, MT4-R5 cells were seeded in 96-well plates and transduced with a library of ISG-encoding SCRPSY vectors (one ISG per well) containing 527 unique human ISG open reading frames. 48 hours after transduction, cells were split into two new plates and infected with a GFP-encoding virus. For single-cycle infections with NHG, 100 µg/ml dextran sulphate was added to the cells at 16 hours post-infection to limit viral spread and cells were fixed at 48 hours post-infection in formaldehyde. For multi-cycle infections using the CH058 GIN viruses, cells were fixed at 96 hours post-infection in formaldehyde. Fixed cells were analysed with flow cytometry using a Guava EasyCyte system (Luminex). Subsequent validation screens of ISG ‘hits’ and candidate effectors were conducted with independent lentiviral preps using the same methods. For the exclusions screens, MT4 and TMZR5 cells were transduced with the genes in Fig 2G as before and 96 hours post-transduction the supernatants were harvested to measure toxicity of the expressed ISGs using the CytoTox-Glow kit (Promega). In a separate experiment, MT4 cells were transduced with the same genes and 96 hours post-transduction the cells were fixed in 4% formaldehyde and stained using the LIVE/DEAD(tm) Fixable Red Dead Cell Stain Kit (Invitrogen) to assess viability of the transduced cells. In a similar fashion MT4-ISRE-GFP cells were transduced and at 96 hours post-transduction cells were fixed in 4% formaldehyde to measure ISRE induction (GFP-positive cells) as surrogate for IFN induction.

### Virus infections and titrations

Suspension cells were seeded immediately prior to infection or treatment. For experiments involving IFN treatment, IFNα14 produced as described above was added 24 hours prior to infection. Azidothymidine (3485) and Nelfinavir (4621) were obtained from the NIH AIDS Reagents Program (catalogue numbers indicated in parentheses) and added 2 hours prior to infection using the indicated concentrations. Virus titrations were carried out as previously described [83]. Cells were infected with a titrated challenge of serially diluted virus-containing supernatant. Cell lines were treated with polyanionic dextran sulfate 17-18 hours post-infection to limit infection to a single cycle (where single cycle infection is indicated). At 48 h after virus challenge, levels of infection were determined via flow cytometry, for either GFP-encoding viruses or GFP-reporter TMZR5 infected cells. The titres plotted are the mean of triplicate (n=3) estimations of the titre extrapolated from different doses within the linear range (error bars represent the standard deviation). For spreading assays, cells were infected with a dose of HIV-1 that resulted in 0.01% of cells GFP+ 24 h post infection. The virus inoculum required for this experiment was calculated based on the number of single-cycle infectious units determined in TMZR5 indicator cells. Cells were sampled every 24 h, fixed, and the levels of infection determined by flow cytometry. Experiments were conducted in quadruplicate (n=4). A typical result from at least two independent experiments is shown.

### Infectious yield assays

TMZR5 cells were seeded in 6- well plates and treated with increasing doses of IFNα14. 24 hours after IFN treatment, cells were challenged with HIV-1 at an MOI of 0.5. At 6 hours post infection, cells were washed once with PBS and pelleted by centrifugation. Supernatant containing inoculum was removed and fresh medium containing the appropriate dose of IFN was used to resuspend the cell pellet and transferred into a fresh 6-well plate. At 46 to 48 hpi, supernatant containing virus was harvested and filtered using a 0.45-m filter and infectivity of the virus was estimated via titration.

### Western blot analyses

For preparation of cell lysates, cell pellets were resuspended in protein sample buffer (12.5% glycerol, 175 mM Tris-HCl [pH 8.5], 2.5% SDS, 70 mM 2-mercaptoethanol, 0.5% bromophenol blue). Proteins were subsequently separated on NuPage 4% to 12% Bis-Tris polyacrylamide gels and transferred onto nitrocellulose membranes. Blots were probed with either anti-actin (JLA20 hybridoma; courtesy of the Developmental Studies Hybridoma Bank, University of Iowa), anti-CD38, anti-CD80, anti FNDC3B, anti-SCARB2 (25284-1-AP, 14292-1-AP, 22605-1-AP, 27102-1-AP; Proteintech) anti-MICB (VPA00747; Bio-Rad) or anti-TMEM140 (SAB1304546; Sigma) primary antibodies. Thereafter, membranes were probed with fluorescently labelled goat anti-rabbit or goat anti-mouse secondary antibodies (Thermo Scientific) and scanned using a LiCor Odyssey scanner.

### Simulations

To illustrate the effect of small differences in growth rate on growth curves generated in the presence of a growth-inhibiting substance, we simulated a logistic growth process for two viruses:

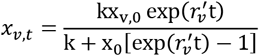

where *x*_*v,t*_ is the number of cells infected by virus *v* at time *t, x*_*v*,0_ is the initial number of infected cells (here fixed to 100), and *k* is the carrying capacity (fixed to 10 000 in all simulations). Finally, 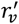 is the effective or realized growth rate of virus *v*, calculated as described below. Viruses were assumed to be growing independently (i.e., in separate wells). To allow different growth rates, the growth rate of virus two was scaled relative to that of virus one by a factor *s*:

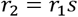

In all simulations, *r*_1_ was held constant at 3, broadly similar to the growth rate measured for CH058 TF (Fig 5D). Similarly, the level of growth rate inhibition, *i*, was allowed to vary between viruses by a scaling factor *c*:

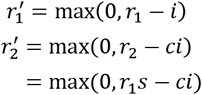

In the first set of simulations, viruses differed in growth rate only, with the second virus having a lower growth rate than the first. This was achieved by varying *s* from 0.6 to 0.95 (i.e., virus two’s growth rate was scaled to between 60% and 95% of virus one’s growth rate) while the inhibition scaling factor *c* was fixed at 1, giving both viruses equal sensitivity to the growth rate inhibitor. In the second set of simulations, the underlying growth rates of both viruses were equal (*s* = 1), but virus 2 was more sensitive to the growth rate inhibitor (*c* > 1). In these simulations, the scaling factor *c* was varied from 1.1 (virus two is 10% more sensitive than virus one) to 1.8 (virus two is 80% more sensitive). In both sets of simulations, the level of inhibition, *i*, was varied such that *r*_1_ would be reduced by between 0 and 90% (Fig 4).

### Analysis of HIV-1 growth rate

To test whether the observed differences in growth curves were the result of growth rate differences, differences in sensitivity to IFN, or both, the spreading assays above were repeated. A maximal dose of 0.5 pg/µL was chosen, as in the initial IFN spreading assays this dose enabled a clear difference between the TF/CC pair with minimal IFN-associated toxicity (∼80% live cells). The remainder of doses were spread at 0.1 pg/µL intervals to capture incremental differences in growth rate.

The data from these assays were modelled as a logistic growth process:

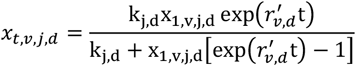

where *x*_*t,v,j,d*_ is the number cells infected at time *t* by virus *v*, in replicate *j* of a given treatment with IFN dose *d*, and *x*_1,*v,j,d*_ is the initial number of infected cells in this replicate (as measured at the first timepoint, 24 hours post inoculation). To account for IFN-toxicity to cells at higher doses, the maximum number of cells available to be infected (i.e. the carrying capacity, *k*_*j*_) was modelled as a function of IFN dose (*d*):

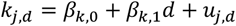

where *β*_*k*,0_ is the mean carrying capacity when no IFN is present, *β*_*k*,1_ is the effect of 1 pg/µl IFN, and *u*_*j,d*_ is a random effect allowing variation in the number of cells available between different replicates of a given treatment.

In the most complex model fitted (here termed the differential sensitivity model), the achieved growth rate of each virus, *r′*_*v,d*_, was modelled as a function of IFN dose, a virus-specific adjustment allowing growth rates to vary between viruses, and an additional virus-specific adjustment for interferon-sensitivity:

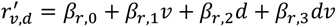

Here, *v* = 0 for the TF virus and 1 for the CC virus. As a result, *β*_*r*,0_ is the growth rate of the TF virus in the absence of IFN (here termed the baseline growth rate), while *β*_*r*,1_ is the adjustment needed to achieve the baseline growth rate of the CC virus. Finally, *β*_*r*,2_ measures the baseline effect of 1 pg/µl IFN on the growth rates of both viruses, while *β*_*r*,3_ allows the CC virus to be more or less sensitive to a given IFN dose than the TF virus. The fit of this model was compared to one without the additional virus-specific adjustment for interferon-sensitivity (i.e. without the *β*_*r*,3_*dv* term), here named the constant sensitivity model.

Models were fit by maximum likelihood using version 3.1-149 of the nlme library in R version 4.0.2 [84, 85]. Confidence intervals for all parameter estimates were generated by re-fitting models to 1000 hierarchical bootstrap samples of the data. For each IFN dose, the available data were truncated as soon as growth curves declined by more than 30% relative to the previous timepoint, with models fit to the remaining data only. This was needed to accommodate the long timescale of these experiments, where both the accumulation of dead cells due to virus infection and release, and the toxicity effects of long-term culture in the presence of IFN, results in a reduction in viable cells that can be infected (Fig 5B). The sensitivity of models to this exclusion was assessed by evaluating a range of cut-off points (including no data removal). Truncation affected primarily the estimated carrying capacity and associated effect sizes (*β*_*k*,0_ and *β*_*k*,1_), with carrying capacity under-estimated when the declining parts of growth curves were included. All other parameter estimates remained broadly similar with overlapping confidence intervals, regardless of the cut-off used, and the differential sensitivity model remained unsupported.

### HIV-1 plasmid sequencing and assembly

40 ng of each plasmid DNA was sheared into approximately 350 base pair in length by sonication using a Covaris Sonicator LE220 (Covaris). Fragmented DNA was uniquely index tagged with NEBNext Multiplex Oligos for Illumina (New England Bio-Labs, E7780S and ES7600S). The Kapa LTP Library Preparation Kit (KAPA Biosystems, Roche7961880001) was deployed in this process. Libraries were quantified and quality controlled with Qubit dsDNA HS kit (ThermoFisher) and Agilent 4200 Tapestation System (Agilent). Equimolar amounts of each library were pooled together and sequenced on the Illumina MiSeq platform using MiSeq Reagent Micro Kit v2 (2x 150-cycles). Plasmid sequences were assembled using SPAdes v3.10.1 with multiple k-mer sizes. Minimum depth of 100 reads and Phred quality of 30 were used for consensus calling of the assembled sequences.

### Analysis of HIV-1 sequences

Using a procedure outlined in [83] to determine the frequency of each amino acid, the Los Alamos National Database (http://www.hiv.lanl.gov/) was used to download all gene sequences available ranging from 4568 sequences for tat to 19237 for nef in subtype B and from 1548 sequence in rev to 4345 in env for subtype C. Only one sequence was selected per patient. Following a codon alignment of each gene, the frequency of amino acids was determined for sites that are different between the paired TF and CC sequences.

## ACKNOWLEDGMENTS

We thank Beatrice Hahn, Stuart Neil, Stephen Goodbourn, the NIH AIDS Reagent Program, and the Developmental Studies Hybridoma Bank at the University of Iowa for reagents, viruses, and cell lines. Schematic of the ISG screening pipeline used in Fig 2E created with BioRender.com.

This study was supported by the Medical Research Council awards MR/P022642/1 (to SJW and SJR), MC_UU_12014/12 (to DLR, NM, JH, VBS); Wellcome Trust awards 201366/Z/16/Z (to SJR) and 217221/Z/19/Z (to NM); Medical Research Council [MC_UU_12018/12] (to AdSF, LT).

---

**S1.**
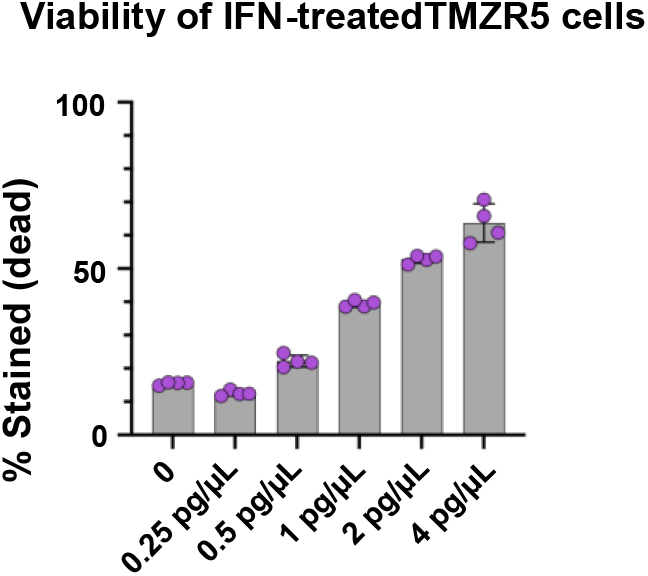
Because of the proapoptotic effect of IFNs, the viability of IFN-treated TMZR5 cells was also assessed in parallel cultures to hose used in Fig 1. Viability was tested using flow cytometry using the LIVE/DEAD fixable dead cell stain kit (Invitrogen)

**S2.**
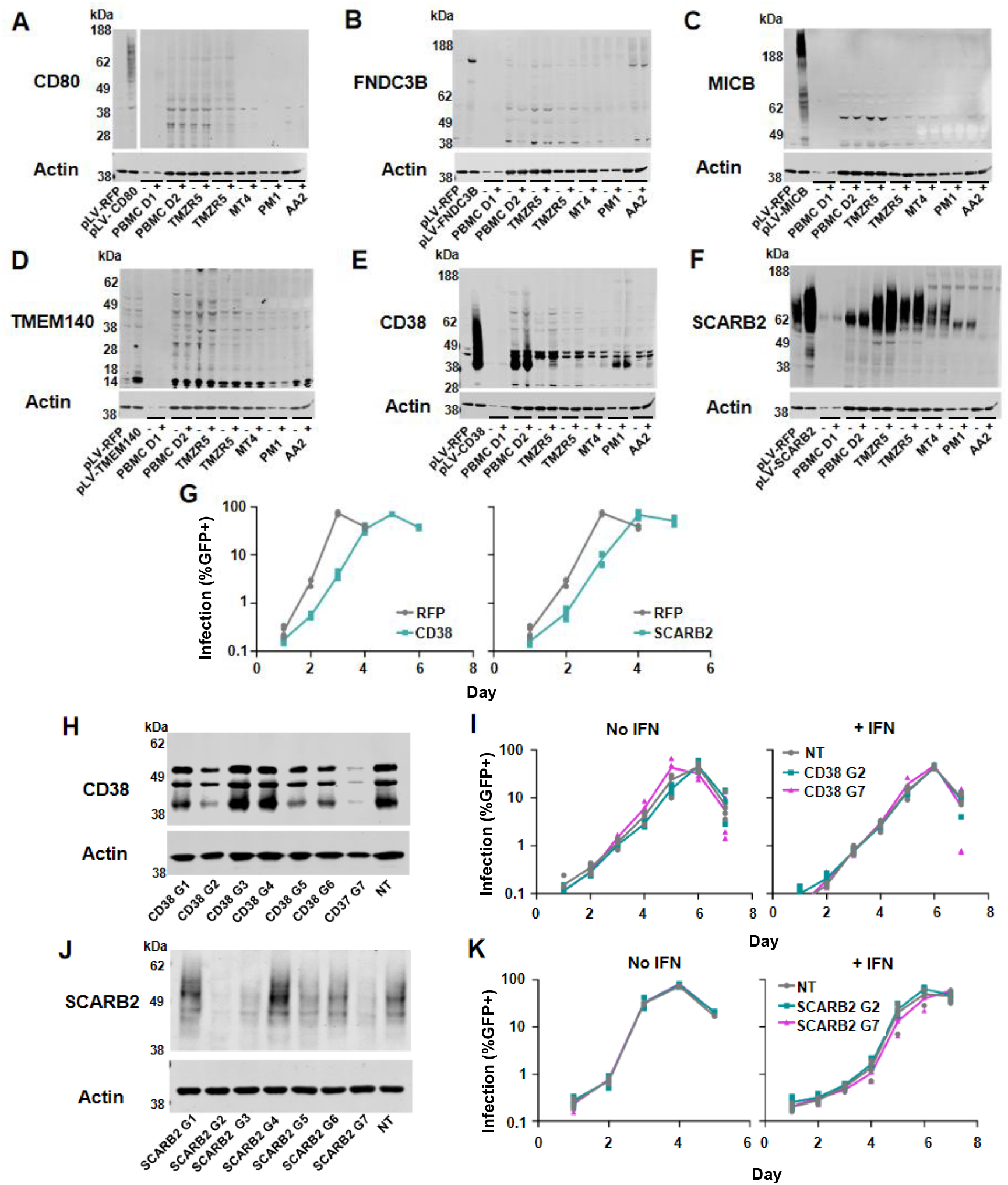
Western blot gels assessing protein expression levels and IFN induction of expression of (A) CD80, (B) FNDC3B, (C) MICB (D) TMEM140, (E) CD38, (F) SCARB2. (G) TMZR5 cells (modified to express CD38 and SCARB2) were challenged with NHG and sampled daily to monitor virus spread. GFP-positive cells were enumerated via flow cytometry. (H) Western blots of the seven CRISPR guides and non-targeting control guide cell lines for CD38 in PM1 cells (I) PM1 cell lines were pre-treated for 24 hours with the IFNα14 doses indicated and were subsequently challenged with NHG and sampled daily to monitor virus spread. (J) Western blots of the seven CRISPR guides and non-targeting control guide cell lines for SCARB2 in TMZR5 cells. (K) TMZR5 cell lines were pre-treated for 24 hours with the IFNα14 doses indicated and were subsequently challenged with NHG and sampled daily to monitor virus spread. Viral spreading replication experiments took place on two occasions, a typical result is shown.

**S3.**
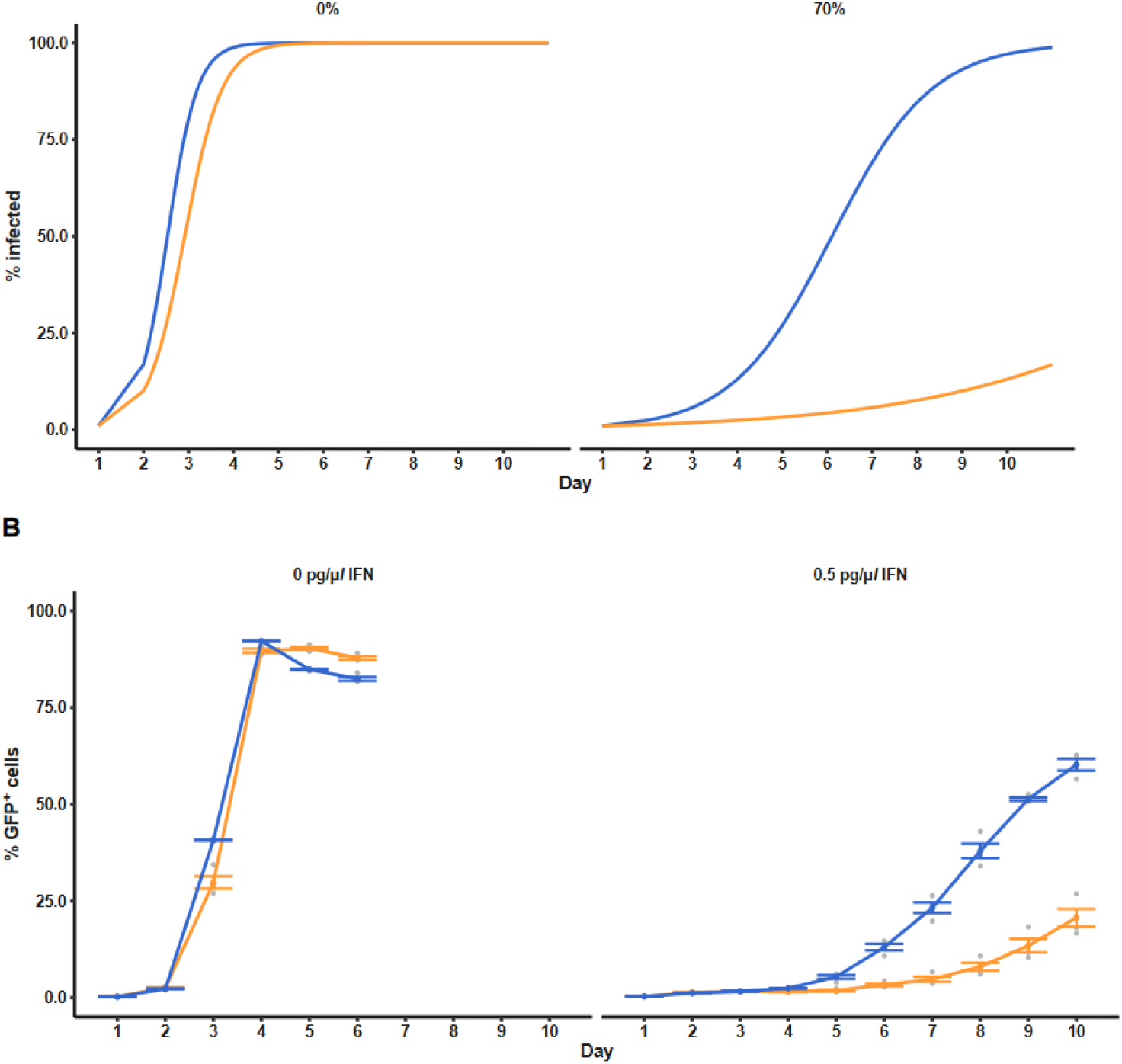
Small growth rate differences become discernible under growth rate inhibition because the lag phase becomes observable. (A) Logistic growth simulation of two viruses, where the growth rate of virus two (orange) is scaled to 0.8 times that of virus one (blue). Both viruses experience the same amount of inhibition; labels above each plot indicate percent inhibition relative to the growth rate of virus 1. (B) Observed growth dynamics with and without interferon. Coloured points show means (+/-standard error) across 4 xperimental replicates, while grey points show individual observations (data as in figure 5; blue: CH085 TF, orange: CH085 CC).

**S4.**
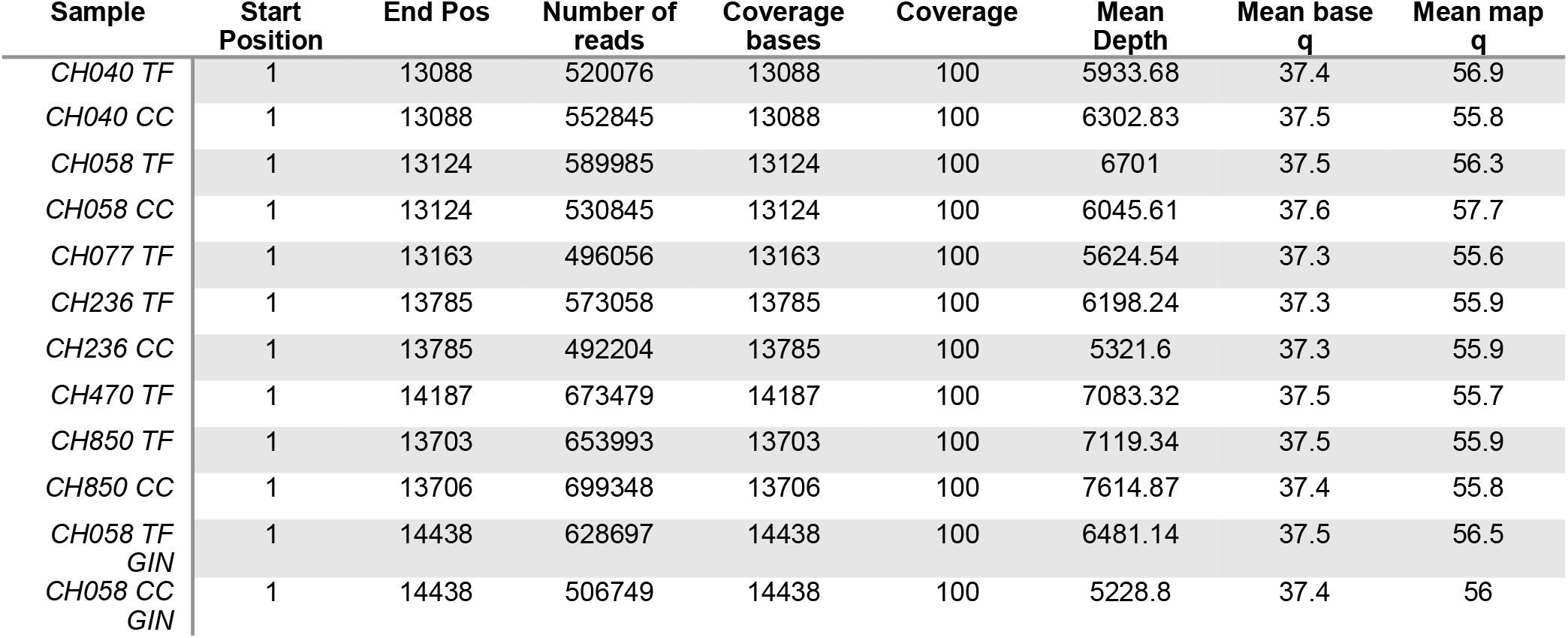
Sequencing coverage statistics

**S5.**
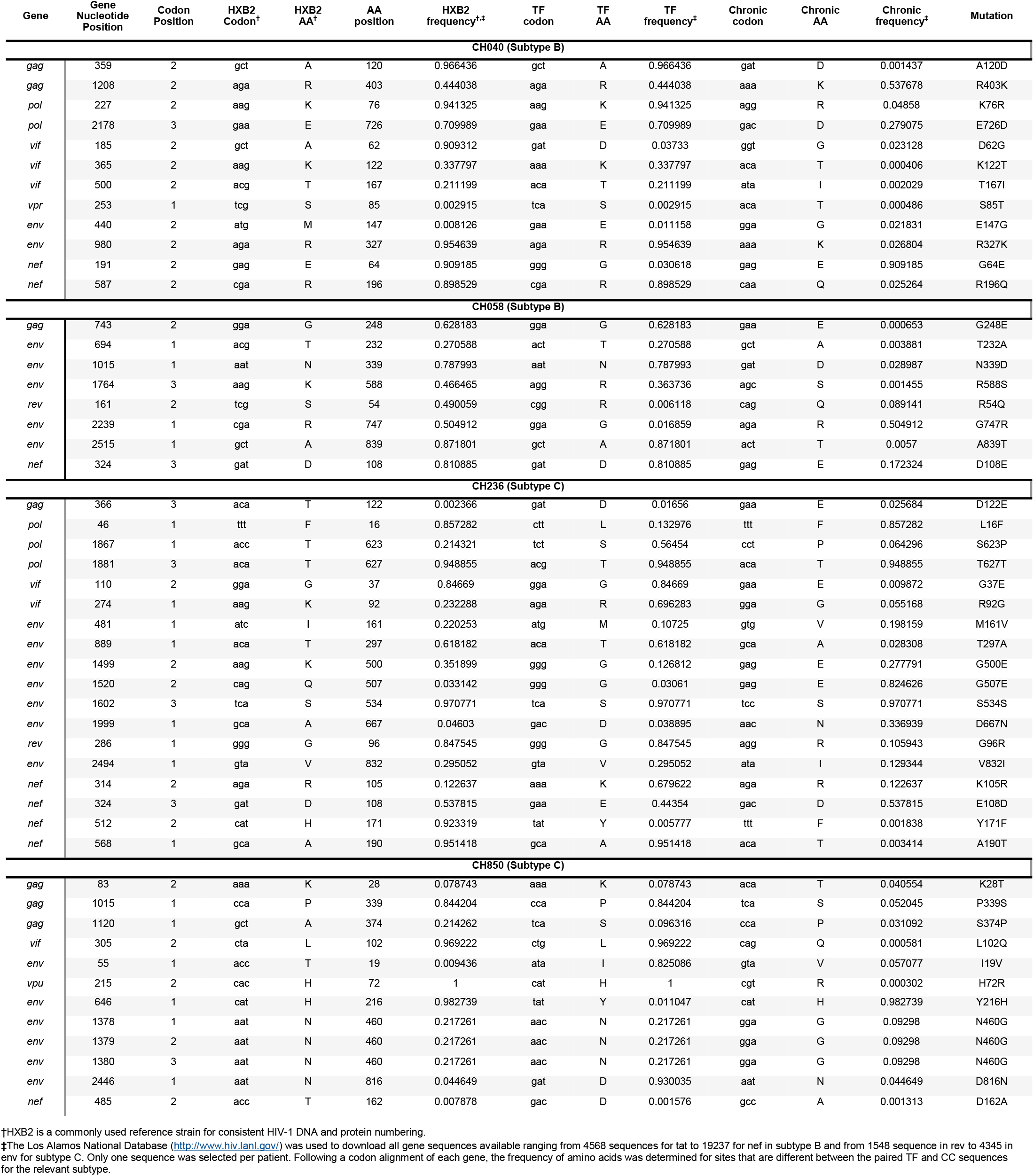
Amino acid frequencies at sites that exhibit amino acid changes between sequenced TF/CC pairs are shown and compared to HxB2 (reference sequence)

**Western blot images used in Fig 2**

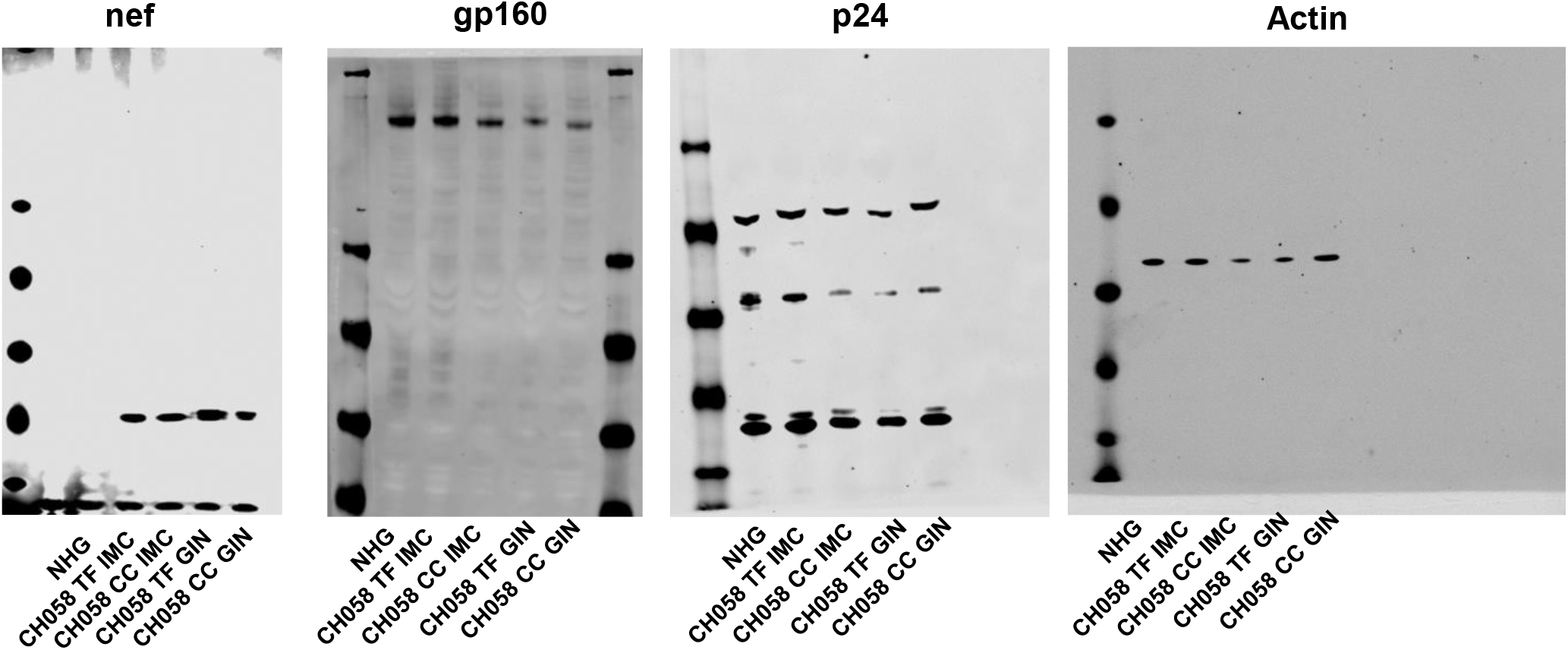

**Western blot files used in S2**

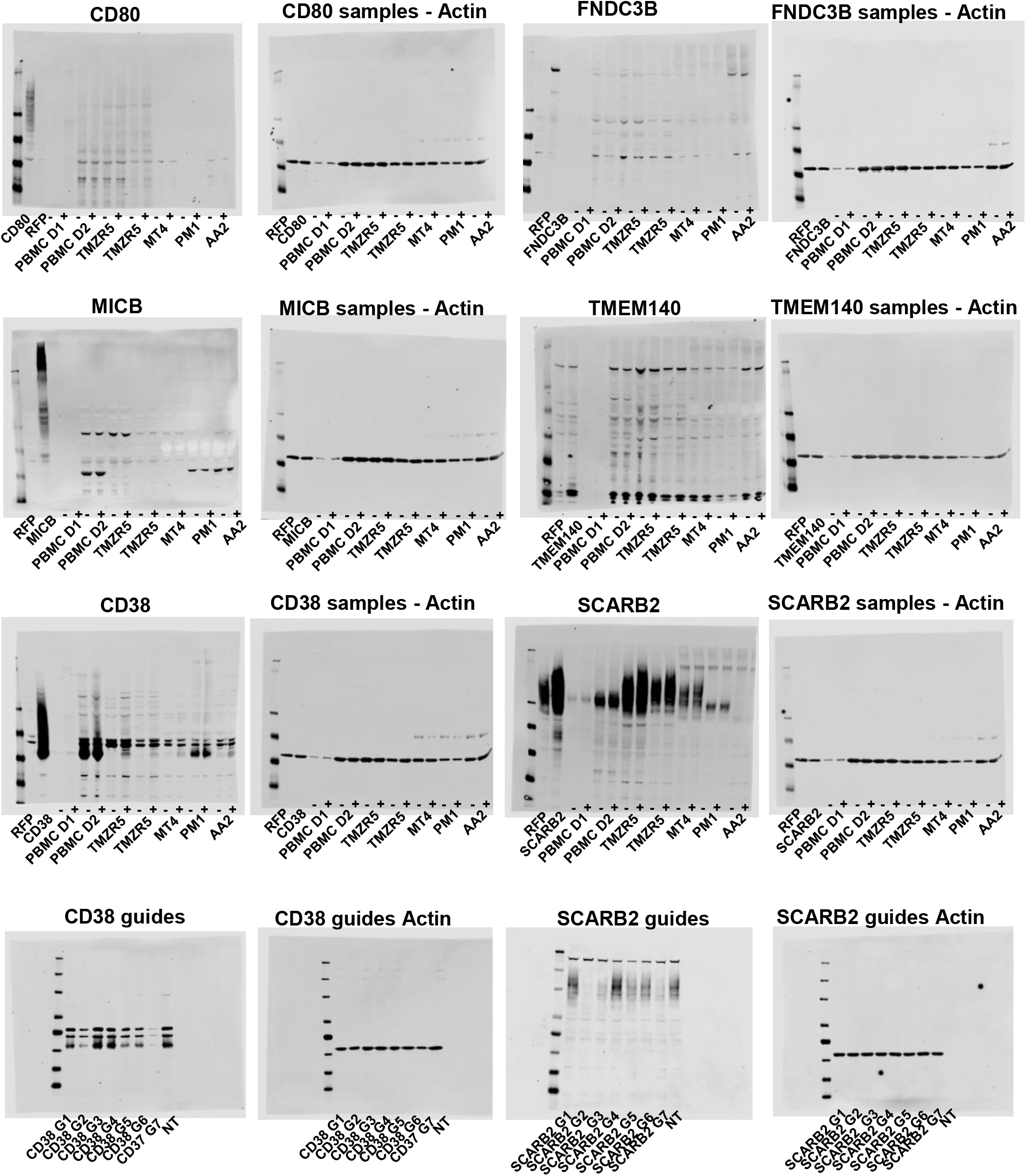

## REFERENCES

1. Stacey AR, Norris PJ, Qin L, Haygreen EA, Taylor E, Heitman J, et al. Induction of a striking systemic cytokine cascade prior to peak viremia in acute human immunodeficiency virus type 1 infection, in contrast to more modest and delayed responses in acute hepatitis B and C virus infections. Journal of virology. 2009;83(8):3719–33.

2. Kane M, Zang TM, Rihn SJ, Zhang F, Kueck T, Alim M, et al. Identification of interferon-stimulated genes with antiretroviral activity. Cell host & microbe. 2016;20(3):392–405.

3. Schoggins JW, Wilson SJ, Panis M, Murphy MY, Jones CT, Bieniasz P, et al. A diverse range of gene products are effectors of the type I interferon antiviral response. Nature. 2011;472(7344):481–5.

4. Royce RA, Sena A, Cates Jr W, Cohen MS. Sexual transmission of HIV. New England Journal of Medicine. 1997;336(15):1072–8.

5. Patel P, Borkowf CB, Brooks JT, Lasry A, Lansky A, Mermin J. Estimating per-act HIV transmission risk: a systematic review. AIDS (London, England). 2014;28(10):1509.

6. Salazar-Gonzalez JF, Bailes E, Pham KT, Salazar MG, Guffey MB, Keele BF, et al. Deciphering human immunodeficiency virus type 1 transmission and early envelope diversification by single-genome amplification and sequencing. Journal of virology. 2008;82(8):3952–70.

7. Keele BF, Giorgi EE, Salazar-Gonzalez JF, Decker JM, Pham KT, Salazar MG, et al. Identification and characterization of transmitted and early founder virus envelopes in primary HIV-1 infection. Proceedings of the National Academy of Sciences. 2008;105(21):7552–7.

8. Abrahams M-R, Anderson JA, Giorgi E, Seoighe C, Mlisana K, Ping L-H, et al. Quantitating the multiplicity of infection with human immunodeficiency virus type 1 subtype C reveals a non-poisson distribution of transmitted variants. Journal of virology. 2009;83(8):3556–67.

9. Villabona-Arenas CJ, Hall M, Lythgoe KA, Gaffney SG, Regoes RR, Hué S, et al. Number of HIV-1 founder variants is determined by the recency of the source partner infection. Science. 2020;369(6499):103-

10. Shaw GM, Hunter E. HIV transmission. Cold Spring Harbor perspectives in medicine. 2012;2(11):a006965.

11. Cohen OJ, Kinter A, Fauci AS. Host factors in the pathogenesis of HIV disease. Immunological reviews. 1997;159(1):31–48.

12. Song H, Pavlicek JW, Cai F, Bhattacharya T, Li H, Iyer SS, et al. Impact of immune escape mutations on HIV-1 fitness in the context of the cognate transmitted/founder genome. Retrovirology. 2012;9(1):1-

13. Haaland RE, Hawkins PA, Salazar-Gonzalez J, Johnson A, Tichacek A, Karita E, et al. Inflammatory genital infections mitigate a severe genetic bottleneck in heterosexual transmission of subtype A and C HIV-1. PLoS Pathog. 2009;5(1):e1000274.

14. Carlson JM, Schaefer M, Monaco DC, Batorsky R, Claiborne DT, Prince J, et al. Selection bias at the heterosexual HIV-1 transmission bottleneck. Science. 2014;345(6193).

15. Joseph SB, Swanstrom R, Kashuba AD, Cohen MS. Bottlenecks in HIV-1 transmission: insights from the study of founder viruses. Nature Reviews Microbiology. 2015;13(7):414–25.

16. Wilen CB, Parrish NF, Pfaff JM, Decker JM, Henning EA, Haim H, et al. Phenotypic and immunologic comparison of clade B transmitted/founder and chronic HIV-1 envelope glycoproteins. Journal of virology. 2011;85(17):8514–27.

17. Gnanakaran S, Bhattacharya T, Daniels M, Keele BF, Hraber PT, Lapedes AS, et al. Recurrent signature patterns in HIV-1 B clade envelope glycoproteins associated with either early or chronic infections. PLoS pathogens. 2011;7(9):e1002209.

18. Derdeyn CA, Decker JM, Bibollet-Ruche F, Mokili JL, Muldoon M, Denham SA, et al. Envelope-constrained neutralization-sensitive HIV-1 after heterosexual transmission. Science. 2004;303(5666):2019–22.

19. Parker ZF, Iyer SS, Wilen CB, Parrish NF, Chikere KC, Lee F-H, et al. Transmitted/founder and chronic HIV-1 envelope proteins are distinguished by differential utilization of CCR5. Journal of virology. 2013;87(5):2401–11.

20. Fenton-May AE, Dibben O, Emmerich T, Ding H, Pfafferott K, Aasa-Chapman MM, et al. Relative resistance of HIV-1 founder viruses to control by interferon-alpha. Retrovirology. 2013;10(1):1–18.

21. Gondim MV, Sherrill-Mix S, Bibollet-Ruche F, Russell RM, Trimboli S, Smith AG, et al. Heightened resistance to host type 1 interferons characterizes HIV-1 at transmission and after antiretroviral therapy interruption. Science Translational Medicine. 2021;13(576).

22. Parrish NF, Gao F, Li H, Giorgi EE, Barbian HJ, Parrish EH, et al. Phenotypic properties of transmitted founder HIV-1. Proceedings of the National Academy of Sciences. 2013;110(17):6626–33.

23. Kmiec D, Iyer SS, Stürzel CM, Sauter D, Hahn BH, Kirchhoff F. Vpu-mediated counteraction of tetherin is a major determinant of HIV-1 interferon resistance. MBio. 2016;7(4):e00934–16.

24. Neil SJ, Zang T, Bieniasz PD. Tetherin inhibits retrovirus release and is antagonized by HIV-1 Vpu. Nature. 2008;451(7177):425–30.

25. Sheehy AM, Gaddis NC, Malim MH. The antiretroviral enzyme APOBEC3G is degraded by the proteasome in response to HIV-1 Vif. Nature medicine. 2003;9(11):1404–7.

26. Mlcochova P, Apolonia L, Kluge SF, Sridharan A, Kirchhoff F, Malim MH, et al. Immune evasion activities of accessory proteins Vpu, Nef and Vif are conserved in acute and chronic HIV-1 infection. Virology. 2015;482:72–8.

27. Oberle CS, Joos B, Rusert P, Campbell NK, Beauparlant D, Kuster H, et al. Tracing HIV-1 transmission: envelope traits of HIV-1 transmitter and recipient pairs. Retrovirology. 2016;13(1):1–20.

28. Deymier MJ, Ende Z, Fenton-May AE, Dilernia DA, Kilembe W, Allen SA, et al. Heterosexual transmission of subtype C HIV-1 selects consensus-like variants without increased replicative capacity or interferon-α resistance. PLoS Pathog. 2015;11(9):e1005154.

29. Iyer SS, Bibollet-Ruche F, Sherrill-Mix S, Learn GH, Plenderleith L, Smith AG, et al. Resistance to type 1 interferons is a major determinant of HIV-1 transmission fitness. Proceedings of the National Academy of Sciences. 2017;114(4):E590–E9.

30. Claiborne DT, Prince JL, Scully E, Macharia G, Micci L, Lawson B, et al. Replicative fitness of transmitted HIV-1 drives acute immune activation, proviral load in memory CD4+ T cells, and disease progression. Proceedings of the National Academy of Sciences. 2015;112(12):E1480–E9.

31. Foster TL, Wilson H, Iyer SS, Coss K, Doores K, Smith S, et al. Resistance of transmitted founder HIV-1 to IFITM-mediated restriction. Cell host & microbe. 2016;20(4):429–42.

32. Wang C, Liu D, Zuo T, Hora B, Cai F, Ding H, et al. Accumulated mutations by 6 months of infection collectively render transmitted/founder HIV-1 significantly less fit. Journal of Infection. 2020;80(2):210-

33. Busnadiego I, Kane M, Rihn SJ, Preugschas HF, Hughes J, Blanco-Melo D, et al. Host and viral determinants of Mx2 antiretroviral activity. Journal of virology. 2014;88(14):7738–52.

34. Stremlau M, Owens CM, Perron MJ, Kiessling M, Autissier P, Sodroski J. The cytoplasmic body component TRIM5α restricts HIV-1 infection in Old World monkeys. Nature. 2004;427(6977):848–53.

35. Kane M, Yadav SS, Bitzegeio J, Kutluay SB, Zang T, Wilson SJ, et al. MX2 is an interferon-induced inhibitor of HIV-1 infection. Nature. 2013;502(7472):563-

36. Compton AA, Bruel T, Porrot F, Mallet A, Sachse M, Euvrard M, et al. IFITM proteins incorporated into HIV-1 virions impair viral fusion and spread. Cell host & microbe. 2014;16(6):736–47.

37. Laguette N, Sobhian B, Casartelli N, Ringeard M, Chable-Bessia C, Ségéral E, et al. SAMHD1 is the dendritic-and myeloid-cell-specific HIV-1 restriction factor counteracted by Vpx. Nature. 2011;474(7353):654–7.

38. Sheehy AM, Gaddis NC, Choi JD, Malim MH. Isolation of a human gene that inhibits HIV-1 infection and is suppressed by the viral Vif protein. Nature. 2002;418(6898):646–50.

39. Li M, Kao E, Gao X, Sandig H, Limmer K, Pavon-Eternod M, et al. Codon-usage-based inhibition of HIV protein synthesis by human schlafen 11. Nature. 2012;491(7422):125–8.

40. Feng J, Wickenhagen A, Turnbull ML, Rezelj VV, Kreher F, Tilston-Lunel NL, et al. Interferon-stimulated gene (ISG)-expression screening reveals the specific antibunyaviral activity of ISG20. Journal of virology. 2018;92(13).

41. Rihn SJ, Aziz MA, Stewart DG, Hughes J, Turnbull ML, Varela M, et al. TRIM69 inhibits vesicular stomatitis Indiana virus. Journal of virology. 2019;93(20).

42. OhAinle M, Helms L, Vermeire J, Roesch F, Humes D, Basom R, et al. A virus-packageable CRISPR screen identifies host factors mediating interferon inhibition of HIV. Elife. 2018;7:e39823.

43. Amara A, Vidy A, Boulla G, Mollier K, Garcia-Perez J, Alcamí J, et al. G protein-dependent CCR5 signaling is not required for efficient infection of primary T lymphocytes and macrophages by R5 human immunodeficiency virus type 1 isolates. Journal of virology. 2003;77(4):2550–8.

44. Zhang Y-j, Hatziioannou T, Zang T, Braaten D, Luban J, Goff SP, et al. Envelope-dependent, cyclophilin-independent effects of glycosaminoglycans on human immunodeficiency virus type 1 attachment and infection. Journal of virology. 2002;76(12):6332–43.

45. Goujon C, Moncorgé O, Bauby H, Doyle T, Ward CC, Schaller T, et al. Human MX2 is an interferon-induced post-entry inhibitor of HIV-1 infection. Nature. 2013;502(7472):559–62.

46. Rusinova I, Forster S, Yu S, Kannan A, Masse M, Cumming H, et al. Interferome v2. 0: an updated database of annotated interferon-regulated genes. Nucleic acids research. 2012;41(D1):D1040–D6.

47. Lu J, Pan Q, Rong L, Liu S-L, Liang C. The IFITM proteins inhibit HIV-1 infection. Journal of virology. 2011;85(5):2126–37.

48. Tartour K, Appourchaux R, Gaillard J, Nguyen X-N, Durand S, Turpin J, et al. IFITM proteins are incorporated onto HIV-1 virion particles and negatively imprint their infectivity. Retrovirology. 2014;11(1):1–14.

49. Perelson AS, Kirschner DE, De Boer R. Dynamics of HIV infection of CD4+ T cells. Mathematical biosciences. 1993;114(1):81–125.

50. Perelson AS, Nelson PW. Mathematical analysis of HIV-1 dynamics in vivo. SIAM review. 1999;41(1):3–44.

51. Ferguson AL, Mann JK, Omarjee S, Ndung’u T, Walker BD, Chakraborty AK. Translating HIV sequences into quantitative fitness landscapes predicts viral vulnerabilities for rational immunogen design. Immunity. 2013;38(3):606–17.

52. Mann JK, Barton JP, Ferguson AL, Omarjee S, Walker BD, Chakraborty A, et al. The fitness landscape of HIV-1 gag: advanced modeling approaches and validation of model predictions by in vitro testing. PLoS Comput Biol. 2014;10(8):e1003776.

53. Rihn SJ, Wilson SJ, Loman NJ, Alim M, Bakker SE, Bhella D, et al. Extreme genetic fragility of the HIV-1 capsid. PLoS Pathog. 2013;9(6):e1003461.

54. Davey NE, Satagopam VP, Santiago-Mozos S, Villacorta-Martin C, Bharat TA, Schneider R, et al. The HIV mutation browser: a resource for human immunodeficiency virus mutagenesis and polymorphism data. PLoS computational biology. 2014;10(12):e1003951.

55. Crawford H, Matthews PC, Schaefer M, Carlson JM, Leslie A, Kilembe W, et al. The hypervariable HIV-1 capsid protein residues comprise HLA-driven CD8+ T-cell escape mutations and covarying HLA-independent polymorphisms. Journal of virology. 2011;85(3):1384–90.

56. Brockman MA, Schneidewind A, Lahaie M, Schmidt A, Miura T, DeSouza I, et al. Escape and compensation from early HLA-B57-mediated cytotoxic T-lymphocyte pressure on human immunodeficiency virus type 1 Gag alter capsid interactions with cyclophilin A. Journal of virology. 2007;81(22):12608–18.

57. Schouest B, Weiler AM, Janaka SK, Myers TA, Das A, Wilder SC, et al. Maintenance of AP-2-dependent functional activities of Nef restricts pathways of immune escape from CD8 T lymphocyte responses. Journal of virology. 2018;92(5):e01822–17.

58. Scanlan CN, Pantophlet R, Wormald MR, Ollmann Saphire E, Stanfield R, Wilson IA, et al. The broadly neutralizing anti-human immunodeficiency virus type 1 antibody 2G12 recognizes a cluster of α1→ 2 mannose residues on the outer face of gp120. Journal of virology. 2002;76(14):7306–21.

59. Alexandre KB, Moore PL, Nonyane M, Gray ES, Ranchobe N, Chakauya E, et al. Mechanisms of HIV-1 subtype C resistance to GRFT, CV-N and SVN. Virology. 2013;446(1-2):66-76.

60. Leitman EM, Willberg CB, Tsai M-H, Chen H, Buus S, Chen F, et al. HLA-B* 4: 02-restricted Env-specific CD8+ T-cell activity has highly potent antiviral efficacy associated with immune control of HIV infection. Journal of virology. 2017;91(22):e00544–17.

61. Waheed AA, Ablan SD, Sowder RC, Roser JD, Schaffner CP, Chertova E, et al. Effect of mutations in the human immunodeficiency virus type 1 protease on cleavage of the gp41 cytoplasmic tail. Journal of virology. 2010;84(6):3121–6.

62. Goonetilleke N, Liu MK, Salazar-Gonzalez JF, Ferrari G, Giorgi E, Ganusov VV, et al. The first T cell response to transmitted/founder virus contributes to the control of acute viremia in HIV-1 infection. Journal of experimental medicine. 2009;206(6):1253–72.

63. Cohen GB, Rangan VS, Chen BK, Smith S, Baltimore D. The human thioesterase II protein binds to a site on HIV-1 Nef critical for CD4 down-regulation. Journal of Biological Chemistry. 2000;275(30):23097-

64. Lundquist CA, Tobiume M, Zhou J, Unutmaz D, Aiken C. Nef-mediated downregulation of CD4 enhances human immunodeficiency virus type 1 replication in primary T lymphocytes. Journal of virology. 2002;76(9):4625–33.

65. Ashokkumar M, Sonawane A, Sperk M, Tripathy SP, Neogi U, Hanna LE. In vitro replicative fitness of early Transmitted founder HIV-1 variants and sensitivity to Interferon alpha. Scientific Reports. 2020;10(1):1–12.

66. Moore CB, John M, James IR, Christiansen FT, Witt CS, Mallal SA. Evidence of HIV-1 adaptation to HLA-restricted immune responses at a population level. Science. 2002;296(5572):1439–43.

67. Allen TM, Altfeld M, Geer SC, Kalife ET, Moore C, O’sullivan KM, et al. Selective escape from CD8+ T-cell responses represents a major driving force of human immunodeficiency virus type 1 (HIV-1) sequence diversity and reveals constraints on HIV-1 evolution. Journal of virology. 2005;79(21):13239–49.

68. Liu Z, Pan Q, Ding S, Qian J, Xu F, Zhou J, et al. The interferon-inducible MxB protein inhibits HIV-1 infection. Cell host & microbe. 2013;14(4):398–410.

69. Wei X, Decker JM, Wang S, Hui H, Kappes JC, Wu X, et al. Antibody neutralization and escape by HIV-1. Nature. 2003;422(6929):307–12.

70. Young D, Andrejeva L, Livingstone A, Goodbourn S, Lamb R, Collins P, et al. Virus replication in engineered human cells that do not respond to interferons. Journal of virology. 2003;77(3):2174–81.

71. Young D, Galiano M, Lemon K, Chen Y-H, Andrejeva J, Duprex W, et al. Mumps virus Enders strain is sensitive to interferon (IFN) despite encoding a functional IFN antagonist. The Journal of general virology. 2009;90(Pt 11):2731.

72. Randall RE, Goodbourn S. Interferons and viruses: an interplay between induction, signalling, antiviral responses and virus countermeasures. Journal of general virology. 2008;89(1):1–47.

73. Rihn SJ, Wilson SJ, Loman NJ, Alim M, Bakker SE, Bhella D, et al. Extreme genetic fragility of the HIV-1 capsid. PLoS pathogens. 2013;9(6):e1003461.

74. Troyer RM, McNevin J, Liu Y, Zhang SC, Krizan RW, Abraha A, et al. Variable fitness impact of HIV-1 escape mutations to cytotoxic T lymphocyte (CTL) response. PLoS Pathog. 2009;5(4):e1000365.

75. Hertoghs N, Nijmeijer BM, van Teijlingen NH, Fenton-May AE, Kaptein TM, van Hamme JL, et al. Sexually transmitted founder HIV-1 viruses are relatively resistant to Langerhans cell-mediated restriction. Plos one. 2019;14(12):e0226651.

76. Sheldon J, Beach NM, Moreno E, Gallego I, Piñeiro D, Martínez-Salas E, et al. Increased replicative fitness can lead to decreased drug sensitivity of hepatitis C virus. Journal of virology. 2014;88(20):12098–111.

77. Gallego I, Sheldon J, Moreno E, Gregori J, Quer J, Esteban JI, et al. Barrier-independent, fitness-associated differences in sofosbuvir efficacy against hepatitis C virus. Antimicrobial agents and chemotherapy. 2016;60(6):3786–93.

78. Gallego I, Gregori J, Soria ME, García-Crespo C, García-Álvarez M, Gómez-González A, et al. Resistance of high fitness hepatitis C virus to lethal mutagenesis. Virology. 2018;523:100–9.

79. Sandler NG, Bosinger SE, Estes JD, Zhu RT, Tharp GK, Boritz E, et al. Type I interferon responses in rhesus macaques prevent SIV infection and slow disease progression. Nature. 2014;511(7511):601–5.

80. Sanjana NE, Shalem O, Zhang F. Improved vectors and genome-wide libraries for CRISPR screening. Nature methods. 2014;11(8):783.

81. Wilson SJ, Schoggins JW, Zang T, Kutluay SB, Jouvenet N, Alim MA, et al. Inhibition of HIV-1 particle assembly by 2′, 3′-cyclic-nucleotide 3′-phosphodiesterase. Cell host & microbe. 2012;12(4):585–97.

82. Ochsenbauer C, Edmonds TG, Ding H, Keele BF, Decker J, Salazar MG, et al. Generation of transmitted/founder HIV-1 infectious molecular clones and characterization of their replication capacity in CD4 T lymphocytes and monocyte-derived macrophages. Journal of virology. 2012;86(5):2715–28.

83. Rihn SJ, Foster TL, Busnadiego I, Aziz MA, Hughes J, Neil SJ, et al. The envelope gene of transmitted HIV-1 resists a late interferon gamma-induced block. Journal of virology. 2017;91(7).

84. Team RC. R: A Language and Environment for Statistical Computing. R Foundation for Statistical Computing; 2020.

85. Pinheiro J BD, DebRoy S, Sarkar D, R Core Team. nlme: Linear and Nonlinear Mixed Effects Models.. R package version 3.1-149 ed2020.

